# IRE1 activity regulates tumour and microenvironment cell lineage states while stratifying localised and metastatic prostate cancer

**DOI:** 10.1101/2025.03.13.643042

**Authors:** D. Doultsinos, I. Tomljanovic, E. Pilalis, S. Abusamra, R Parmentier, I.E. Bridges, S. Figiel, Y. Zekri, J. Hester, D. Leach, C. Bevan, A.D. Lamb, W. Zwart, A. Chatziioannou, C. Le Magnen, I.G. Mills

## Abstract

Prostate cancer (PCa) is an androgen receptor (AR) driven, high-incidence disease significantly contributing to cancer mortality. PCa is in need of better risk stratification at diagnosis and treatment outcomes in patients at high risk of metastasis. The unfolded protein response (UPR) is an AR-dependent process. However, the impact of the UPR transducer IRE1 on AR-dependent biology and treatment resistance has not been defined. We use diverse pre-clinical models of stress response to describe IRE1 activity impact on multiple disease stages and demonstrate its involvement with poor prognosis (RB1 loss), and cell lineage determination (club phenotypes). Integrating clinical transcriptomic datasets, we chart IRE1 activity throughout PCa evolution by developing a PCa-specific, IRE1 activity gene set (IRE1_18) reflecting both tumoral and micro-environmental niches. IRE1_18 can determine tumoral identity, inform androgen deprivation treatment suitability, prognosticate localised and metastatic disease independently from AR activity, and guide IRE1 modulation as a novel combination therapeutic.

**Graphical Abstract created using Biorender.com:** 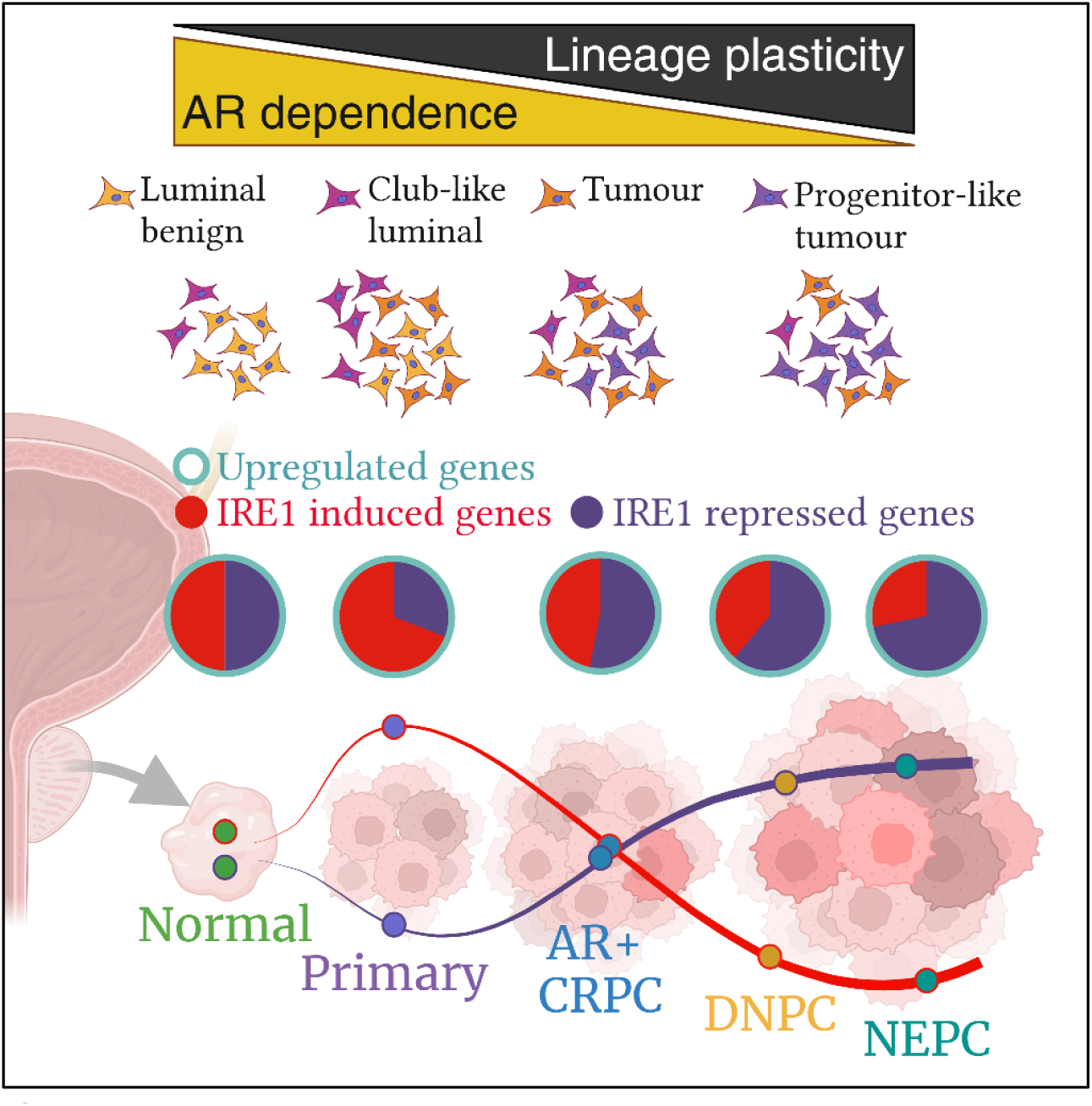

## Introduction

Prostate cancer (PCa) is a heterogenous, multifocal, high-incidence disease that affects one in eight men globally. PCa evolution depends on transcriptional events and epigenetic modifications, forming an adaptive response to intrinsic/extrinsic stresses, culminating in lineage reprogramming and treatment resistance^1^. PCa results in ∼360000 deaths annually^2^ due to metastatic progression, supported by treatment/stress-resistant cancer cells which experience high proliferative rates and androgen receptor (AR)-driven high metabolic demand.

These events require high protein turnover, leading to misfolded protein accumulation, causing Endoplasmic Reticulum (ER) stress. The unfolded protein response (UPR) resolves this stress through three main transducers, Inositol Requiring Enzyme alpha (IRE1α), protein kinase RNA-like endoplasmic reticulum kinase (PERK) and activating transcription factor 6α (ATF6). IRE1α, hereafter referred to as IRE1 and encoded by the gene *ERN1*, is the most evolutionarily conserved transducer of the UPR and signals through two distinct downstream functional outputs^3^. Firstly, through the unconventional splicing of the transcript encoding XBP1, a transcription factor, resulting in a non-canonical splice variant lacking a 60-nucleotide sequence which encodes the active protein (XBP1s). Secondly, through the targeted degradation/decay (RIDD) of a subset of mRNA (e.g., *DGAT2*^4^, *BLOC1S1*^5^) and miRNA transcripts to ease transcriptional load^6^. Both functions require IRE1 endoribonuclease activity and can be thought of as a continuum of early (XBP1s) and late (RIDD) overlapping responses that are designed to adapt to stress.

Using short (up to 24 hour) pre-clinical model treatments we have previously shown that the IRE1-XBP1 axis contains AR-induced genes, notably *ERN1* itself^7^. Targeting IRE1 in androgen-dependent pre-clinical models can restrain tumorigenesis^8^. However, it remains unclear which IRE1-dependent genes or biological processes underpin these responses and how IRE1 activity may change in response to sustained stress, reflecting PCa evolution. Most studies so far have focused on the role of IRE1 as an activator of XBP1 impacting tumour development^9^ largely ignoring IRE1-RIDD impact on RNA stability which could be restraining the emergence of cancers that are more difficult to treat.

In this study we used bulk, single cell (SC) and spatial (ST) RNA sequencing data from two-dimensional (cell lines) and three-dimensional models (patient derived explants (PDEs)), as well as patient biopsy samples to describe how IRE1 activity changes throughout PCa evolution. Crucially, we found that adaptation to IRE1 loss correlated with cell identities usually driven by prognostically poor genomic alterations such as RB1 loss. We also observed that IRE1-deficient cells develop a phenotype resembling club cells, epithelial cells that interact with immunosuppressive microenvironmental cells. We produced a novel IRE1 activity gene set, IRE1_18, representing significant ERN1 transcriptomic perturbations in PCa-specific models. IRE1_18 is a proxy of multiple signalling pathways linked to PCa progression and can chart PCa evolution from early stage to treatment resistant disease while prognosticating large clinical cohorts. Current clinical practices impact therapeutic effectiveness on PCa mortality^10^ and therefore, using IRE1 activity to identify tumours with treatment resistance traits early, could be transformative of patient care. IRE1 activity could be used to stratify patients for therapy intensification, surveillance or suitability of IRE1/UPR modulation as novel combination therapeutics alongside conventional treatments.

## Results

### Changes in AR activity correlate with significant changes in IRE1 activity in PCa cell lines and clinical datasets

To capture steady state adaptations to AR-induced physiological stresses we cultured LNCaP cells in charcoal stripped serum media (CSM) or full serum media (FM) for 72 hours or treated them with the synthetic androgen R1881 or EtOH for 12 hours. Whole transcriptome sequencing of these conditions followed by gene set enrichment analysis (GSEA) revealed biological pathways negatively enriched in CSM vs FM while positively enriched in the R1881 vs EtOH conditions including the androgen response, fatty acid metabolism, MTORC1 signalling, oxidative phosphorylation and cholesterol homeostasis (Figure 1A, B, Supplementary tables S1, S2)). Canonical AR targets reflecting AR activity were significantly upregulated in the R1881 treatment whilst significantly downregulated in the CSM condition (Supplementary Figure S1A, Supplementary tables S3, S4).

**Figure 1.**
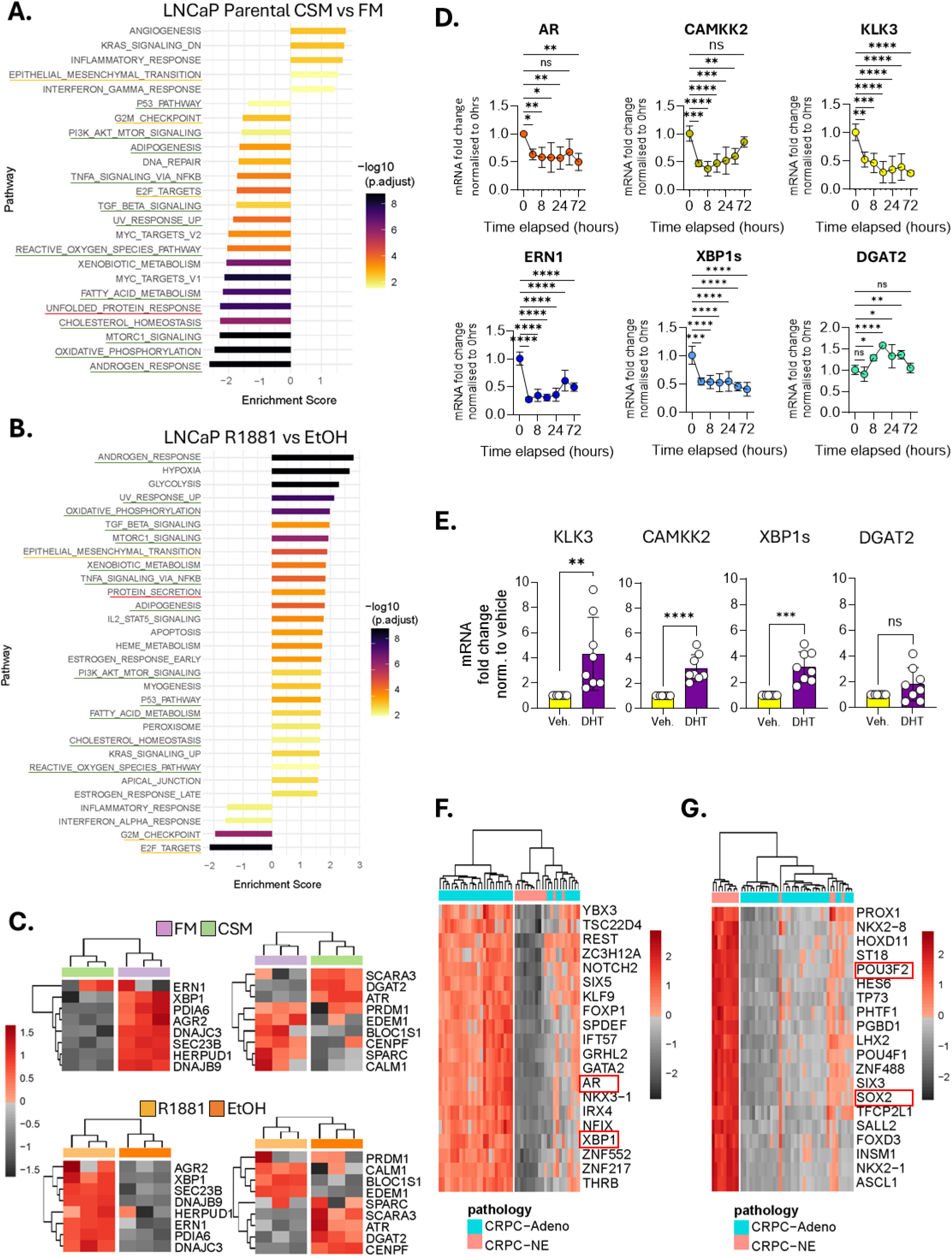
Changes in AR activity correlate with significant changes in IRE1 activity in pre-clinical models and clinical datasets. Barplots displaying GSEA normalized enrichment scores (NES) for significantly enriched MSigDB Hallmark gene sets in: LNCaP cells grown in charcoal stripped media (CSM) for 72 hours compared to LNCaP cells grown in full media (FM) for 72 hours (A), and LNCaP cells treated with R1881 for 12 hours compared to LNCaP cells treated with ethanol (EtOH) for 12 hours (B). Bar length indicates the normalized enrichment score and bar colour signifies the statistical significance. Hallmarks activated in B) but suppressed in A) are underlined in green. Hallmarks activated or suppressed in both A) and B) are underlined in orange. Hallmarks representing protein homeostasis are underlined in red. C) Expression heatmaps of selected “IRE1 induced” (left) and “IRE1 repressed” (right) genes according to the literature in LNCaP cells in CSM (green) vs FM (purple), (top); and R1881 (yellow) vs EtOH (orange) (bottom). Colour scale represents expression z-score from black (lowest) to red (highest). D) mRNA fold change of androgen receptor activity markers (full length AR, CAMKK2, KLK3) and IRE1 activity markers (ERN1, XBP1s, DGAT2) in LNCaP parental cells grown over 72 hours in CSM, normalised to untreated FM control. Statistical analysis was performed using an ordinary one-way ANOVA with Dunnett’s multiple comparisons test and single pooled variance. Four stars signify adjusted p value of <0.0001. E) mRNA fold change of AR and IRE1 activity markers KLK3, CAMKK2, XBP1s and DGAT2 in patient derived explants (PDEs) treated with dihydrotestosterone (DHT), normalised to PDEs treated with vehicle control. Statistical analysis was performed using a T-test per condition. Each star signifies an order of magnitude of significance <0.05. F, G) Heatmaps representing regulons significantly altered in neuroendocrine (NEPC, pink) castration resistant (CRPC) tumours compared to adenocarcinoma CRPC (light blue). Regulon levels are represented as single-sample enrichment scores with individual regulons denoted by the name of their corresponding transcription factor (TF). Significantly activated and suppressed regulons are shown, with AR, XBP1 (AR, IRE1 activity), SOX2 and POU3F2 (reprogramming) in red boxes. Colour scale represents regulon enrichment z-scores from lowest in greyscale to highest in red-scale.

Some Hallmarks were only significant upon androgen treatment or androgen deprivation. These included a positive enrichment of protein secretion and negative enrichment of the Unfolded Protein Response (UPR) in R1881 vs EtOH and CSM vs FM conditions respectively (Figure 1A, B). The UPR Hallmark is not IRE1 specific containing target genes of all UPR transducers (ATF4, ATF6 and XBP1s) while RIDD may not be represented within it. Consequently, we used a curated list of XBP1, and RIDD target genes to correlate alterations in AR signalling with IRE1 activity. Whereas XBP1 target genes were largely androgen-induced, RIDD targets were not consistently androgen dependent (Figure 1C, Supplementary figure S1A).

UPR responses are tissue dependent and therefore, we determined how AR and IRE1 target gene expression changed over 72-hour CSM culture of LNCaP compared to FM. A decrease in IRE1 activity (*XBP1s* downregulated, RIDD targets *DGAT2*, *BLOC1S1* upregulated) was seen as early as 8 hours, mirroring full length AR levels and activity (Figure 1D). Similar profiles were observed at the protein level (Supplementary Figure S1B). Simultaneously we observed that levels of *ATF4* and *eIF2α* either did not significantly or consistently change (Supplementary Figure S1C).

Hallmarks including E2F targets, G2M checkpoint and epithelial mesenchymal transition EMT (Figure 1A) were partially androgen dependent. Therefore, we hypothesised that they may be regulated by other AR-dependent global mechanisms such as the IRE1 axis. We expanded our pre-clinical model repertoire and assessed canonical AR and IRE1 target gene expression in patient derived explants (PDEs)^11^ treated with, or deprived of, dihydrotestosterone (DHT) for 12 hours. AR response markers *KLK3* and *CAMKK2* were significantly upregulated upon DHT treatment. *ERN1* itself and *DGAT2*, *BLOC1S1* were not significantly altered but XBP1s was significantly upregulated (Figure 1E, Supplementary figure S1D).

We then tested if this AR-IRE1 activity correlation could be replicated within a clinical cohort representing AR positive and AR negative tumours. Regulon analyses of 48 castration resistant prostate cancer (CRPC) samples split into CRPC-Adenocarcinoma (n=35, AR positive) versus CRPC-neuroendocrine prostate cancer (NEPC, n=13, AR negative)^12^ showed that AR and XBP1 regulons were both less active in NEPC samples while reprogramming markers pertaining to stem-like lineages such as SOX2 and POU3F2 were increasingly active (Figure 1F, G). As such, transcription factor (TF) activity correlates with IRE1 activity and PCa lineage.

### Knocking out IRE1 induces changes in lineage characteristics of AR dependent PCa cells

Having established that IRE1 activity changes with PCa progression alongside the AR we investigated the adaptation of AR positive cells (LNCaP) to stable IRE1 ablation, using an IRE1 CRISPR-knockout (LNCaP ERN1^-/-^)^13^, lacking IRE1 at steady state or chemical ER stress (tunicamycin/thapsigargin) (Supplementary Figure S2A, B). Bulk RNAseq confirmed multiple canonical “IRE1 induced” and “IRE1 repressed” genes were downregulated and upregulated respectively in ERN1^-/-^ cells (Supplementary figure S2C).

Any observed biology in the ERN1^-/-^ cells could be attributed to the cell line generation selection pressure. To definitively link physiological observations to the absence of IRE1 activity we stably expressed an IRE1 mutant in the ERN1^-/-^ cells carrying a truncation mutation at tryptophan 780^14^ thus lacking the effector ribonuclease domain. These new ERN1^-/-^ Q780* cells were IRE1 activity deficient at steady state and ER stress conditions (Supplementary Figures S2D-G).

We then documented the growth of LNCaP, LNCaP ERN1^-/-^ and LNCaP ERN1^-/-^ Q780* in FM or CSM. ERN1^-/-^ and ERN1^-/-^ Q780* doubling time was 25% and 32% lower than parental LNCaP in FM, respectively. In CSM, ERN1^-/-^ and ERN1^-/-^ Q780* doubling time was increased by 50% compared to FM whilst parental cells stopped growing (Figure 2A) entirely. To document the biological functions underlying this phenotype we performed pathway analysis on whole transcriptome data from these conditions. E2F targets, G2M checkpoint, and the EMT were significantly enriched in ERN1^-/-^ samples (Figure 2B, Supplementary Table S5).

**Figure 2.**
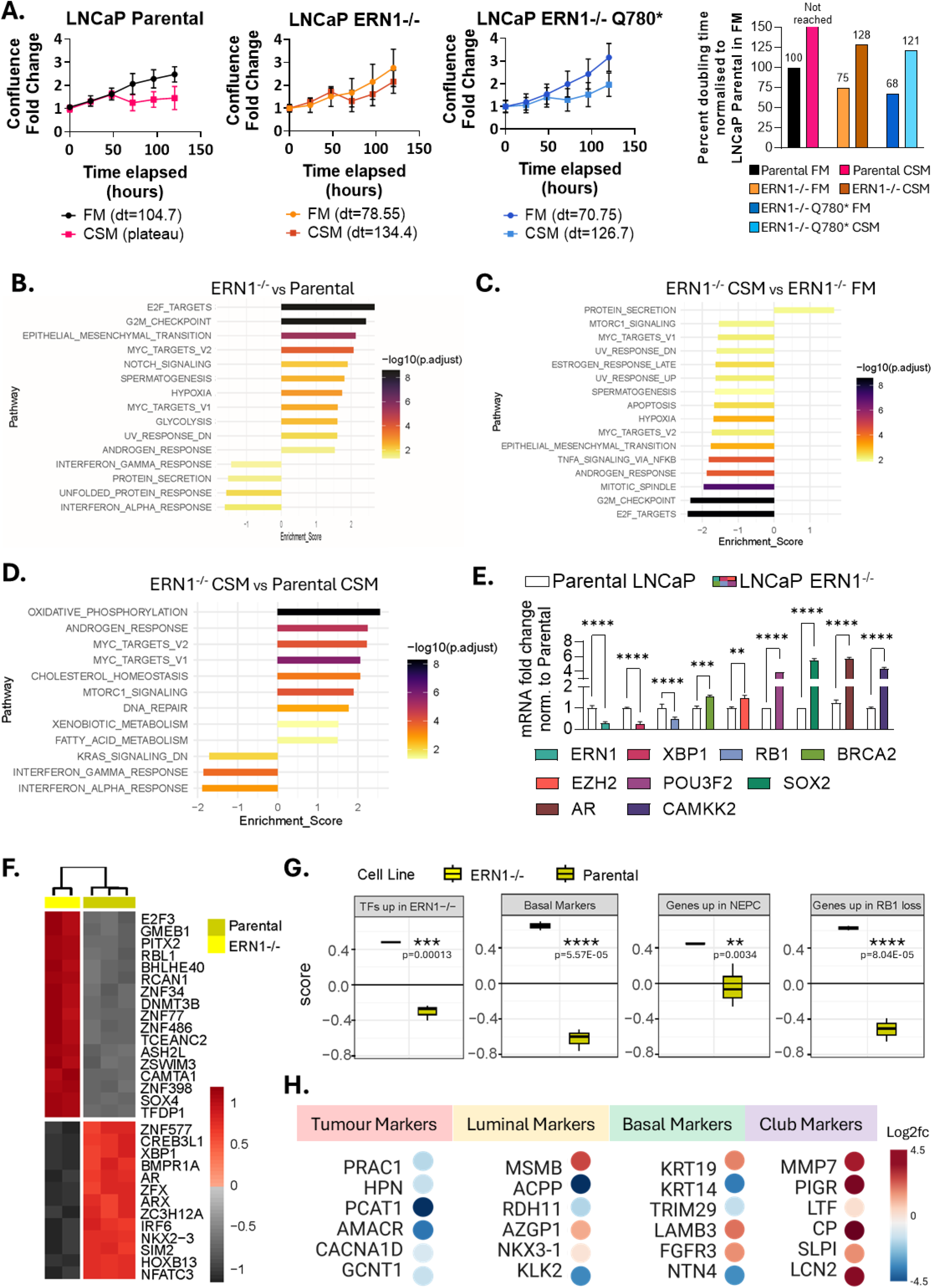
Knocking out IRE1 induces lineage changes in AR-dependent PCa cells. A) Growth curves of LNCaP parental, LNCaP ERN1-/- and LNCaP ERN1-/- Q780* cells grown over 120 hours in FM or CSM. Confluence fold change was normalised to Incucyte readings taken at the 0-hour time point of each condition. Exponential (Malthusian) growth curve equations were used to estimate the doubling time of each cell line in FM or CSM. GSEA normalized enrichment scores (NES) for significantly enriched MSigDB Hallmark gene sets in LNCaP ERN1-/- cells compared to LNCaP parental cells grown in FM (B), LNCaP ERN1-/- cells grown in CSM compared to LNCaP ERN1-/- cells grown in FM (C), and LNCaP ERN1-/- cells compared to LNCaP parental cells grown in CSM (D). Bar length indicates the normalized enrichment score (NES) and bar colour signifies the statistical significance. E) mRNA fold change of ERN1, XBP1, cell cycle markers RB1, BRCA2, and lineage reprogramming markers EZH2, POU3F2, SOX2 in LNCaP ERN1-/- normalized to parental LNCaP. Statistical analysis was performed using an ordinary two-way ANOVA and Sidak’s multiple comparisons test with single pooled variance. Heatmap representing regulons significantly altered in LNCaP ERN1-/- (IKO, light blue) compared to parental LNCaP (pink). Top 30 statistically significant regulons shown (p<0.001). Regulon levels are represented as single-sample enrichment scores with individual regulons denoted by the name of their corresponding transcription factor (TF). Colour scale represents regulon enrichment z-scores (from lowest in greyscale to highest in red-scale). G) GSVA single-sample gene set enrichment scores enrichment scores in LNCaP ERN1-/- cells (IKO, yellow) compared to LNCaP parental cells (mustard) shown from left to right for gene sets including: TFs with regulons activated in ERN1-/- cells, genes upregulated in the RB1-loss signature from Ertel et al., a basal cell signature from Aggarwal et al, and the genes upregulated in the NEPC signature from Beltran et al. Statistical testing was performed using limma. H) Bulk RNAseq log2 fold change expression levels in ERN1^-/-^ cells compared to parental, of genes widely reported to be associated with tumour, luminal epithelial, basal epithelial and club epithelial cells. Downregulated genes in blue, upregulated genes in red. Created using Biorender.com. Each experiment was performed at least in duplicate. Four stars signify adjusted p value of <0.0001.

Consequently, androgen deprivation-induced growth arrest and DNA repair abrogation on PCa cells might be a consequence of both the repression of XBP1 activity and the stabilization of RIDD targets. We thus compared significantly altered Hallmarks induced by androgen deprivation with those altered by knocking out IRE1. ERN1^-/-^ cells were still sensitive to CSM as E2F targets, G2M checkpoint and AR Response were negatively enriched compared to ERN1^-/-^ cells grown in FM (Figure 2C, Supplementary Table S6). However, these Hallmarks were not affected by androgen deprivation to the same degree as DNA repair, AR response and MTORC1 were upregulated in ERN1^-/-^ compared to parental cells grown in CSM (Figure 2D, Supplementary Table S7).

We observed that low *XBP1* correlated with high activity of reprogramming TFs (*SOX2, POU3F2*) in clinical samples (Figure 1F, G) and that IRE1 loss leads to cell cycle de-regulation in PCa cell lines (Figure 2A-D). Consequently, we measured mRNA of lineage reprogramming^15,16^ and cell cycle regulation^17^ genes with known clinical association in PCa. *RB1* was significantly downregulated whilst *BRCA2, SOX2 EZH2* and *POU3F2* mRNA was significantly upregulated in ERN1^-/-^ cells in FM (Figure 2E). Therefore, adaptation to IRE1 activity loss changed the capacity of LNCaP cells to respond to androgen deprivation whilst retaining their AR axis.

We then mapped TF repertoires (regulons) significantly affected by *ERN1* knockout. Stress response (CREB3L1), inflammatory signalling (NFATC3, IRF6), cell cycling (E2F3) and embryogenesis/stemness/differentiation regulons (SOX4, PITX2, HOXB13) were significant. RBL1 was positively enriched (Figure 2F, Supplementary table S8) which has been shown to regulate cell cycle in response to RB1 loss^18^.

Gene signatures associated with RB1 loss, NEPC and basal phenotypes (Supplementary Table S9) segregated ERN1^-/-^ from parental cells, in a similar manner to the regulons (Figure 2G). Thus, knocking out IRE1 in cells that are AR-dependent, and retain some luminal epithelial character, could change their lineage characteristics. Comparing luminal, basal and club cells within a PCa SC dataset^19^ we observed that TFs with regulons activated upon IRE1-loss were significantly upregulated in basal and club cells compared to luminal epithelial cells (Supplementary Figure 3A). We then documented the enrichment of genes upregulated in tumoral, luminal, basal and club cells^20^ (Supplementary Table S10) in the ERN1^-/-^ transcriptome. ERN1^-/-^ cells were positively enriched for club^21–23^ while negatively enriched for tumoral and luminal cell markers, showing decreases of *AMACR*, ACPP and NTN4 but increases in *MMP7, PIGR, MSMB* and *KRT19* (Figure 2H, Supplementary Figure S3B).

### IRE1 perturbation affects cell cycle and lineage reprogramming markers and responses to physiological androgen deprivation therapy (ADT)

Adaptation to IRE1 loss leads to an epithelial club-like lineage shift histologically associated with inflammatory atrophy^21^, that can result from AR blockade^23^. Low IRE1 activity correlates with ADT resistant clinical presentations (Figure 1) and IRE1 deficient cells express high RBL1 and low RB1 RNA levels. Lineage plasticity is facilitated by knocking RB1 out, or knocking RB1 out alongside TP53 in LNCaP cells ^24,25^ which are reported to be resistant to replication stress, have high proliferative rates and increased DNA repair capacity^26^. Intriguingly, Hallmarks affected by RB1 loss or RB1 and TP53 loss were similar to IRE1 loss (Figures 3A, 3B, Supplementary Tables S11, S12) showing upregulation in E2F, G2M checkpoint and MYC targets.

**Figure 3.**
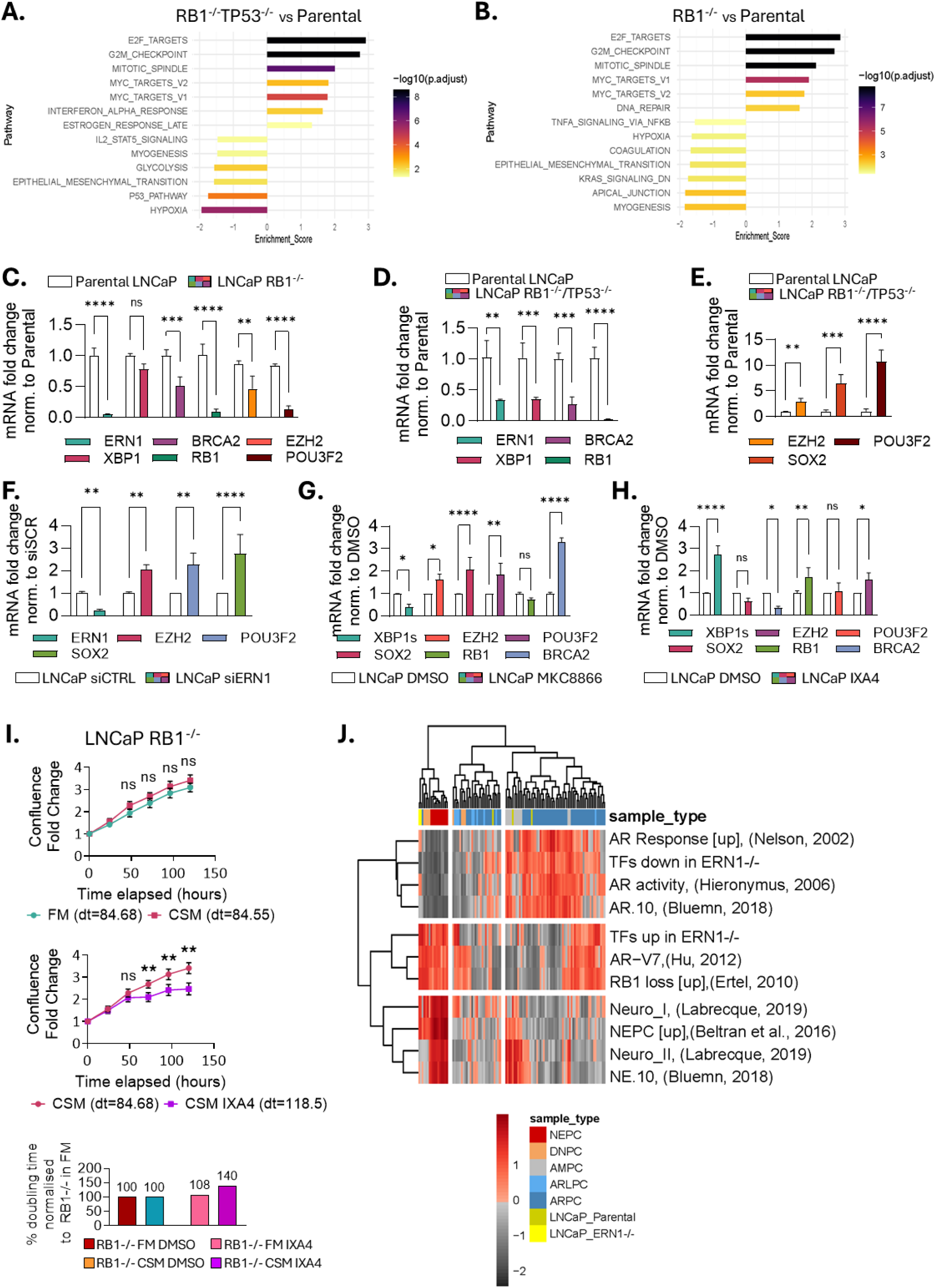
IRE1 activity impacts AR responses in RB1 loss. GSEA normalized enrichment scores (NES) for significantly enriched MSigDB Hallmark (H) gene sets in: LNCaP RB1^-/-^TP53^-/-^ cells compared to LNCaP parental cells grown in FM (A), and LNCaP RB1-/- cells grown in CSM compared to LNCaP parental cells grown in FM (B). Bar length indicates the normalized enrichment score (NES) and bar colour signifies the statistical significance. C, D, E) mRNA fold change of *ERN1, XBP1, RB1, BRCA2, EZH2, POU3F2, SOX2* in C) LNCaP RB1-/- normalised to parental LNCaP and D, E) LNCaP RB1^-/-^TP53^-/-^ cells normalised to parental LNCaP. F) mRNA fold change of *ERN1, EZH2, POU3F2, SOX2* in LNCaP parental cells treated with siRNA against ERN1 normalised to scramble control. mRNA fold change of *XBP1s, EZH2, POU3F2, SOX2, RB1, BRCA2* in LNCaP parental cells treated with G) MKC8866 or H) IXA4 normalised to DMSO. Statistical analysis was performed using an ordinary two-way ANOVA and Sidak’s multiple comparisons test with single pooled variance. I) Growth curves of LNCaP RB1^-/-^ cells grown over 120 hours in FM (green), CSM (red) or in CSM in the presence of subtoxic (25μM) doses of IXA4 (purple). Confluence fold change was normalised to Incucyte readings taken at the 0-hour time point of each condition. T-tests were used for individual time point comparisons between FM and CSM; CSM and CSM plus IXA4. Exponential (Malthusian) growth curve equations were used to estimate the doubling time of each condition. J) Heatmap of GSVA enrichment scores for 11 selected PCa gene signatures shown across samples from the Labrecque patient cohort (N=98) together with LNCaP parental and ERN1-/- cell line samples (N=5). Colour scale represents GSVA enrichment z-scores (from lowest in greyscale to highest in red-scale). Each experiment was performed at least in duplicate. Four stars signify adjusted p value of <0.0001.

We then tested whether IRE1 activity was altered in RB1 deficient cells. *ERN1, BRCA2*, *RB1*, *EZH2* and *POU3F2* were significantly downregulated in RB1^-/-^ cells (Figure 3C). RB1^-/-^/TP53^-/-^ cells had significantly lower levels of both *ERN1*, *XBP1s, BRCA2* and *RB1* (Figure 3D) while higher levels of *EZH2*, *POU3F2* and *SOX2* (Figure 3E) compared to parental.

*ERN1* siRNA silencing significantly upregulated *POU3F2*, *EZH2*, *SOX2* (Figure 3F). Treatment with the salicylaldehyde IRE1 inhibitor MKC8866^27^ caused significant upregulation of *EZH2*, *POU3F2*, *SOX2* and *BRCA2* (Figure 3G) while treatment with the IRE1 activator IXA4^28^, caused significant upregulation of *RB1* and downregulation of *BRCA2* with reprogramming markers showing little change (Figure 3H).

We correlated resistance to ADT with IRE1 activity loss. Therefore, we tested the effect of pharmacological IRE1 activity upregulation on the ability of treatment resistant cells to grow in CSM. CSM did not affect growth of either RB1^-/-^ or RB1^-/-^/TP53^-/-^ LNCaP cells. However, addition of subtoxic doses of the IRE1 activator IXA4, slowed their growth in CSM. RB1^-/-^ doubling time increased by 40% after 24 hours whilst RB1^-/-^/TP53^-/-^ only changed by 10% after 72-hours compared to FM and DMSO (Figure 3I, Supplementary Figure S4A).

Then, in a metastatic, castration resistant PCa (mCRPC, 98 tumours from 55 men)^29^ cohort, we investigated whether the patterns of enrichment of gene signatures comprising TFs with regulons significantly activated or suppressed in ERN1^-/-^ (Figure 2F) could differentiate AR positive from AR negative mCRPC alongside several AR and NEPC gene signatures. The set of TFs with regulons suppressed in ERN1-/- - clustered with AR activity signatures whilst the TFs with regulons activated in ERN1-/- clustered with RB1-loss and ARV-7 activity signatures^30,31^, distinguishing AR naïve from AR agnostic or NEPC/DNPC PCa. Overall, ERN1-/- cells clustered with NEPC and double-negative for NEPC and AR PCa (DNPC) as well as some AR-low (ARLPC) mCRPC samples (Figure 3J). Consequently, IRE1 transcriptomic repertoires may correlate with AR driven and agnostic phenotypes by playing a key role in lineage plasticity, cell cycle and ultimately treatment response.

### Enzalutamide treatment induces a UPR downregulation which may act as a negative feedback loop that leads to treatment resistance to ADT in PCa

Adaptation to IRE1 activity loss may push AR responsive cells towards an ADT resistant phenotype, marking the starting point of IRE1 activity saturation and the shift from an active phenotype to a passive, adapted phenotype. ADT could catalyse this key event in the shift from early stage to late-stage disease progression. Consequently, we investigated the effect of increasing concentrations of Enzalutamide (ENZA) on LNCaP, LNCaP ERN1^-/-^ and LNCaP ERN1^-/-^Q780* cells over 12 hours. Both IRE1-deficient cell lines had significantly higher ENZA IC50s (Figure 4A, Supplementary Figure S4B).

**Figure 4.**
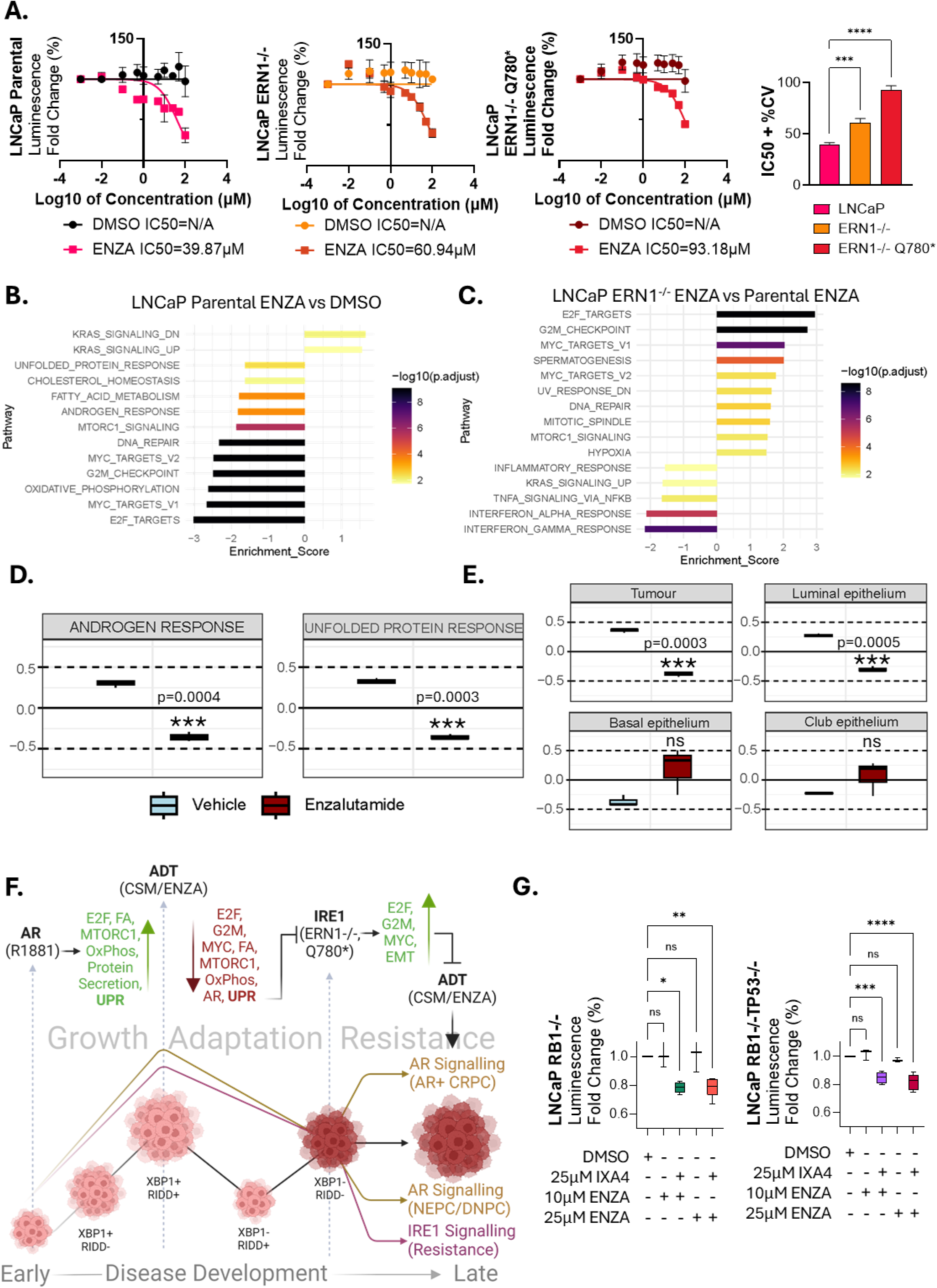
Enzalutamide treatment downregulates UPR responses which contribute to acquired ADT resistance. A) Enzalutamide (ENZA) concentration gradient measuring cytotoxicity through ATP MTT assays in LNCaP parental, LNCaP ERN1^-/-^ and LNCAP ERN1^-/-^ Q780* cells treated for 12 hours with increasing concentrations of ENZA. Logarithmic concentration vs normalised response curves were fit to calculate the IC50 of ENZA for each cell line. Bar chart represents ENZA IC50 comparison between the cell lines. Maximum variance used to calculate significance by ordinary one-way ANOVA with multiple comparisons. B) GSEA normalized enrichment scores (NES) of LNCaP parental cells treated with 10μM ENZA for 12 hours compared to DMSO or C) of LNCaP ERN1^-/-^ cells treated with 10μM ENZA for 12 hours compared to parental LNCaP cells treated with the same conditions. Bar length indicates the normalized enrichment score (NES) and bar colour signifies the statistical significance. Box plots of GSVA enrichment scores for D) Hallmark Androgen Response, Hallmark Unfolded Protein Response and E) gene lists specific for tumour, luminal, club and basal epithelial cells, benchmarked in explants treated with 10μM Enzalutamide (red, right) or vehicle control (blue, left). Statistical testing was performed using limma (lmFit function). F) Schematic summarising observations of figures 1-4. During AR driven tumour growth represented by the R1881 dataset, multiple key mechanisms are upregulated including the UPR. In the eventuality of ADT represented by the CSM and ENZA datasets, these processes are reversed/ downregulated. However, removing IRE1 activity in isolation leads to an increase in processes that directly contradict the effects of ADT leading to partial treatment resistance. Created using Biorender.com. G) Cell titer glo assay, fluorescence fold change of treatment resistant LNCaP RB1^-/-^ and LNCaP RB1^-/-^TP53^-/-^ cells treated with 50μM and 100μM ENZA in the presence or absence of 25μM IXA4. Statistical analysis was performed using an ordinary one-way ANOVA with Dunnett’s multiple comparisons test and single pooled variance. Each experiment was performed at least in duplicate. Four stars signify adjusted p value of <0.0001.

ENZA treatment of parental LNCaP, induced suppression of AR response, DNA repair, MTORC1, E2F targets and the UPR (Figure 4B, Supplementary Table S13). Comparing ERN1-/- to parental LNCaP treated with ENZA, we observed positive significant enrichment for E2F targets, G2M checkpoint and DNA repair (Figure 4C, Supplementary Table S14), corroborating our CSM results (Figures 1,2). In addition, we compared ERN1^-/-^ responses to ENZA, to publicly available RB1^-/-^/TP53^-/-^ LNCaP ENZA response data observing strong similarities between the two. Only inflammatory, hypoxia and MTOR signalling significantly differentiating the responses of these two cell lines to ENZA (Supplementary Figure S4C, Supplementary Tables S15, S16). Having seen that ENZA treatment reduces UPR/IRE1 activity in cell lines we tested its effects on more complex, heterogenous PDEs. ENZA treatment not only suppressed AR and UPR hallmarks (Figure 4D) while increasing inflammatory responses and EMT (Supplemental Figure S4D, Supplementary Table S17, S18), but also significantly decreased luminal and tumoral markers while inducing enrichment trends of club and basal repertoires (Figure 4D, Supplementary Table S10).

Thus, IRE1 activity suppression may play a pivotal role in acquired treatment resistance (Figure 4F) through lineage adaptation to ADT. We then tested whether IRE1 activation could sensitise RB1 deficient cells to ENZA. Whereas the IRE1 activator IXA4 had no effect on viability when used alone (not shown), it sensitised both ENZA resistant (Supplementary Figure S4E, F) RB1^-/-^ and RB1^-/-^TP53^-/-^ cells to physiologically relevant doses of ENZA over a 12-hour treatment (Figure 4G).

Consequently, ENZA treatment suppresses IRE1 activity in cell lines and PDEs, treatment resistant PCa pre-clinical models have low IRE1 activity recapitulating clinical observations and a short-term increase in IRE1 activity enhances responses to AR antagonism.

### Integrative analysis of sequencing datasets gives rise to a novel IRE1 activity gene set (IRE1_18) that represents multiple biological functions that drive PCa dynamic disease progression

We identified PCa-specific, IRE1-responsive genes, labelling them “IRE1 activity induced” (significantly downregulated in ERN1^-/-^) or “IRE1 activity repressed” (significantly upregulated in ERN1^-/-^) (Supplementary Table S19) by integrating results from multiple RNAseq datasets (41 genes, Figure 5A, Supplementary Figure S5A). This gene set represents key PCa progression biological functions comprising cell adhesion, motility, cell cycle, DNA repair, and protein trafficking, while correlating with changes in cell cycle, EMT and metabolism PCa-associated genes (Supplementary Figure S5B, S5C Supplementary Tables S20, S21).

**Figure 5.**
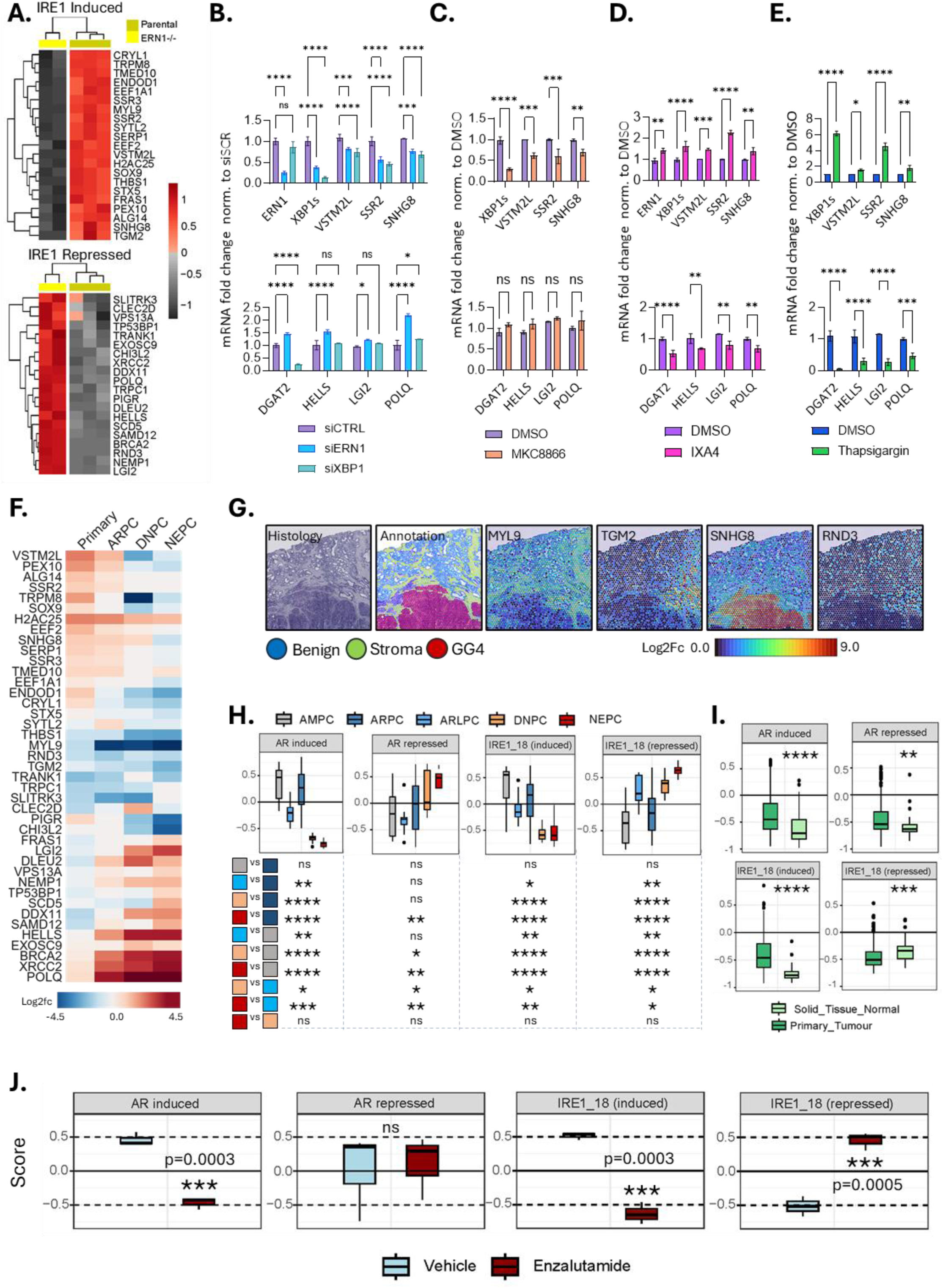
Multiple sequencing datasets give rise to a novel IRE1 activity gene set that represents key PCa biological functions and dynamic disease progression. A) Gene expression heatmaps of genes in the IRE1 activity signature defined as “IRE1 induced” (top) when downregulated in LNCaP ERN1^-/-^ cells (yellow) compared to parental (mustard) and as “IRE1 repressed” (bottom) when upregulated in LNCaP ERN1^-/-^ cells compared to parental. mRNA fold change of *ERN1, XBP1s, DGAT2, VSTM2L, SSR2, SNHG8, HELLS, LGI2* in LNCaP parental cells treated with B) siRNA against ERN1 or XBP1 for 48 hours normalised to scramble control, C) 5μM MKC8866 for 5 hours normalised to DMSO, D) 25μM IXA4 for 5 hours normalised to DMSO and E) 100nM Thapsigargin for 5 hours normalised to DMSO. Statistical analysis was performed using an ordinary two-way ANOVA and Sidak’s multiple comparisons test with single pooled variance. F) Heatmap of the IRE1 activity gene set expression fold change in primary, AR positive CRPC, NEPC and DNPC patient cohorts compared to normal tissue (Prostate Cancer Atlas). Fold changes calculated using “Normal” as control in the DESeq2 Prostate Cancer Atlas function. Downregulated genes in blue, upregulated in red. G) Histological annotations of 6.5 mm^2^ tissue slide where benign (blue), stroma (green) and grade group 4 (GG4) cribiform tumour is represented (red). Loupe browser generated distribution maps of individual genes (*MYL9, TGM2, RND3, SNHG8*) across these histological designations showing either generalised or specific localisation to benign or tumoral tissue. Box plots of GSVA enrichment scores for AR activity and IRE1_18 gene sets, benchmarked in H) mCRPC (Labrecque) and I) localised disease (TCGA PRAD) datasets. Patient populations represented include AR positive (dark blue), AR low (light blue), amphicrine (grey), NEPC (red), double negative (DNPC, orange), normal tissue (light green) and primary tumour (dark green). J) Further testing of AR activity and IRE1_18 gene sets were benchmarked in explants treated with Enzalutamide (10μM, red, right) or vehicle control (blue, left). Statistical testing was performed using limma (lmFit function). Each experiment was performed at least in duplicate. Four stars signify adjusted p value of <0.0001. Heatmap in F) was created using Biorender.com.

We then investigated whether this approach yielded direct IRE1 activity targets within IRE1_41. XBP1 ChIP-seq analysis of LNCaP cells treated with R1881 for 24 hours showed that multiple “IRE1 induced” genes displayed XBP1 peaks directly in their promoter (Supplementary figure S6A). RNA silencing of IRE1 or XBP1 downregulated “IRE1 induced” and upregulated “IRE1 repressed” genes (Figure 5B). The same effect was achieved by MKC8866 (IRE1 RNAse inhibitor) treatment (Figure 5C) while the opposite with IXA4 (Figure 5D) or thapsigargin (Figure 5E) treatment. To ensure that these effects were not cell-line specific, the same analyses were also performed in C4-2B cells and similar effects were observed (Supplementary Figure S6B-F).

We then assessed AR/IRE1 activity canonical genes in a compendium of integrated RNA-seq datasets comprising over 1000 clinical tissue samples ranging from benign to NEPC (Prostate Cancer Atlas^32^). We also applied signatures of basal and luminal epithelial cell lineages to these data^33^. The basal signature expression mirrored AR repression while AR induction signature mirrored both the luminal signature and IRE1 induction (Supplementary Figure S7A). We then investigated the new IRE1 activity gene set in these samples, documenting the pseudo-time disease progression trajectory attributed to the “IRE1 induced” and “IRE1 repressed” components (Supplementary Figure S7B). A particular subgroup of genes (*MYL9, THBS1, TRANK1, TGM2, RND3 and CHI3L2)* was consistently downregulated compared to normal tissue (Figure 5F).

The IRE1 activity gene set was derived from significant differential expression upon IRE1 perturbation in pre-clinical cancer models. However, some of these genes under IRE1 control could be enriched in other cell types. ST^34^ data expression maps encompassing benign, stromal and localised grade group (GG)4 histological annotations, showed that signal was concentrated exclusively in the benign/stromal region; either ubiquitously (*RND3, MYL9*) or in very specific benign sub-compartments (*TGM2*). Some genes were tumour-region specific with the majority of “IRE1 induced” genes (e.g., SNHG8) enriched in GG4 (Figure 5G, Supplementary Figure S7C) while also significantly upregulated in tumoral compared to normal epithelial cells in two separate SC datasets of primary tumour and matched benign samples ^19,35^ (Supplementary Figure S7D). Recapitulating multicellularity, IRE1 activity could distinguish between SC luminal, basal and club cells.

Using these ST and SC sequencing filters, we selected refined IRE1 activity gene sets that identify tumoral and peri-tumoral regions. We applied further redundancy filters (Supplementary Figure S8A) based on expression patterns across clinical designations in the Prostate Cancer Atlas, which segregated AR+ from AR-tumours (*VSTM2L, TRPM8*) or displayed increasing gradient expression (*HELLS, XRCC2, BRCA2*) and decreasing gradient expression patterns (*SNHG8, EEF2*). We thus produced shorter filtered gene sets that were applied to segregating mCRPC (Labrecque) or localised (PRAD TCGA) cohorts (Supplementary Figure S8B, S8C). The final set of 18 genes (labelled IRE1_18) still represented diverse biological functions and cell types (Supplementary Figure S8D, Supplementary Table S22). Both “IRE1 induced” and “IRE1 repressed” parts of IRE1_18 differentiated AR-positive and AR-negative tumours from normal tissue (Supplementary Figure S8E, Supplementary Tables S23, S24).

We next assessed how IRE1_18 clustered metastatic (Labrecque) and localised (TCGA-PRAD) patient cohorts compared to AR signatures. The “IRE1 induced” part of IRE1_18 was significantly reduced in NEPC/DNPC mCRPC samples compared to AR positive mCRPC samples showing similarity to a 10-gene AR induced signature^36^. Within the AR positive mCRPC groups, “IRE1 repressed” subset of IRE1_18 most negatively enriched in the amphicrine group whilst most positively enriched in the AR-low group (Figure 5H). In the TCGA-PRAD, the “IRE1 induced” gene set was significantly decreased in primary disease compared to benign tissue. The exact opposite was observed in the “IRE1 repressed” gene set (Figure 5I). Encouragingly, both parts of IRE1_18 segregated AR-positive from AR-negative tumours and primary tumours from benign tissue in a manner comparable to AR activity signatures.

Since IRE1_18 performed well in segregating benign from tumoral as well as metastatic CRPC samples characterised according to AR status we determined whether IRE1_18 could distinguish PDEs treated with ENZA versus ones treated with vehicle control. The IRE1_18 induced component was significantly downregulated in ENZA treated samples showing similarities to canonical AR activity (AR induced) signatures. The IRE1_18 repressed component conversely, was significantly upregulated, in ENZA treated PDEs while the AR repressed signature showed no change compared to vehicle control.

As such, the IRE1_18 score comprising some direct IRE1 targets shows that IRE1 activity is a proxy of multiple tissue types and as such can segregate benign from tumoral tissue and AR positive from AR negative metastatic CRPC as well as indicate ENZA response in multicellular, heterogenous PDEs.

### Novel IRE1 activity gene set distinguishes between histological grades, between benign and tumour-bearing virtual biopsies and correlates with multiple clinical attributes

IRE1_18 is a proxy of multiple biological functions, cell types and treatment responses and identifies or prognosticates both localised and metastatic disease. Recently, we described how gene expression profiling of virtual tumour and benign biopsies reveals differences between grade groups and tumour clones not reflected in bulk analyses. Therefore, bulk analyses of whole biopsies or tumour-only areas, as used in clinical practice, may provide inaccurate assessments^37^ as tissue type, tumour grade and clonal composition all influence gene expression.

We then tested if IRE1_18 could be confounded by heterogeneity. Pseudo bulk analysis of tumour bearing virtual biopsy using IRE1_18 showed transcriptomic changes attributed to both tumoral or peri-tumoral tissue. Upregulation of *CHI3L2*, *TGM2* and *TRPM8*, *SNHG8*, *VSTM2L* (Supplementary Figure S9A, B) reflected peri-tumoral (*TGM2*, *CHI3L2*) or tumoral (*SNHG8*, *TRPM8, VSTM2L*) regions (Supplementary Figure S9C). We proceeded to determine if IRE1_18 could identify high grade tumour, profile a bulk virtual biopsy and place this profile in an exact subgroup of patients within a larger external clinical cohort. IRE1_18 genes separating GG4 from local benign in a 6.5mm^34^, showed the same pattern in a virtual biopsy containing both GG2 and GG4. Since this patient solely underwent radical prostatectomy, we expected their virtual biopsy to correspond to primary disease and indeed, the expression profile of IRE1_18 corresponds to the primary disease subgroup of the Prostate Cancer Atlas (Figure 6A).

**Figure 6.**
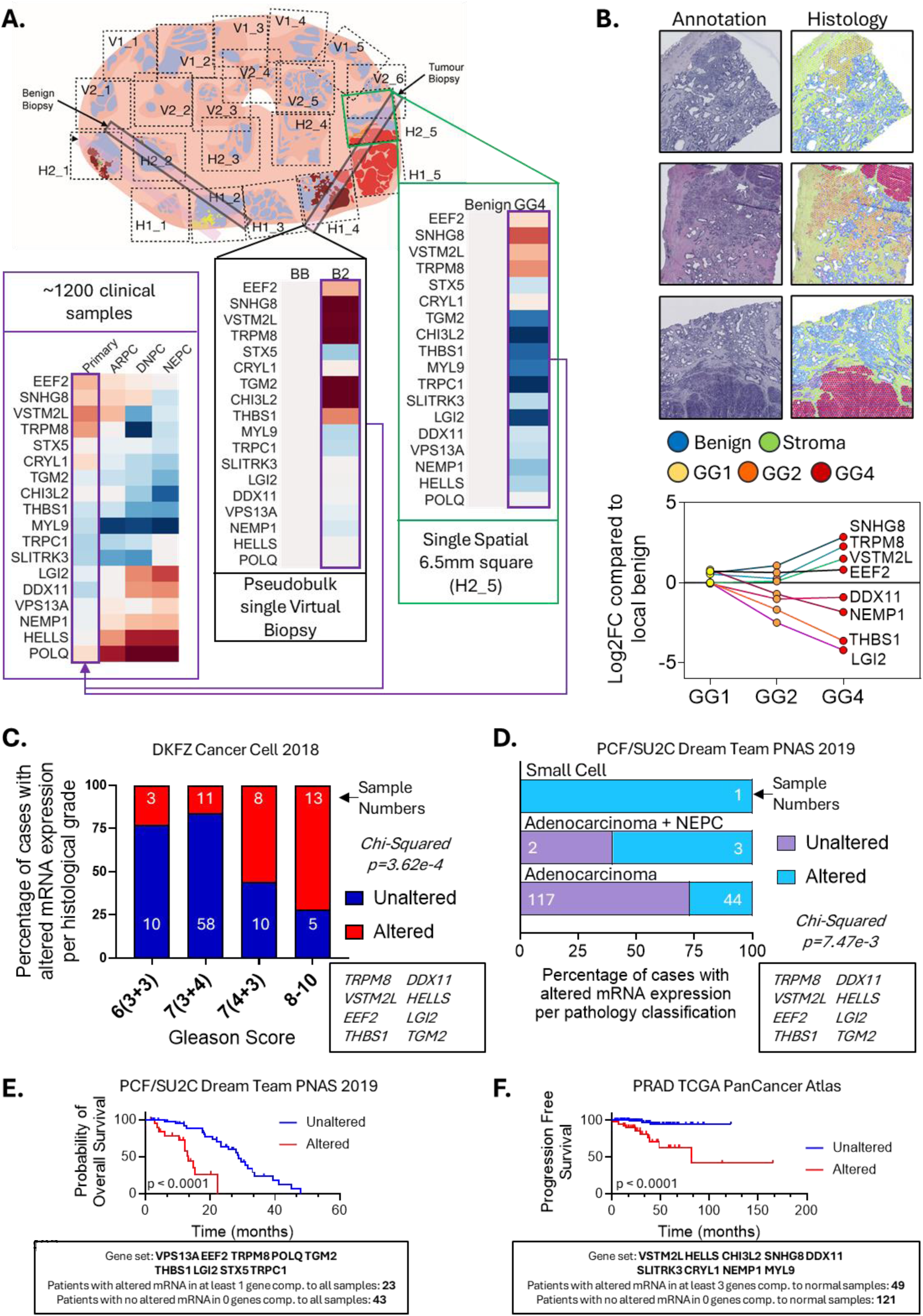
IRE1 activity may act as proxy of multiple clinical attributes. A) Schematic of whole radical prostatectomy axial disc with coded dashed squares signifying corresponding spatial transcriptomic 6.5mm2 squares. As previously described three rectangles show virtual biopsy placements corresponding to completely benign or tumour bearing 1.2mm x 18 mm standard transurethral (TRUS) sampling. From right to left, log fold change expression of the IRE1 activity signature genes is represented in heatmaps locally comparing GG4 to benign; comparing a benign virtual biopsy to a tumour bearing virtual biopsy carrying GG2 and GG4; and comparing primary, AR positive CRPC, DNPC and NEPC bulk samples to normal tissue. B) Histological annotations (left) from three separate 6.5 mm^2^ squares from the same radical prostatectomy representing multifocality and heterogeneity. Grade groups (GG) 1 (yellow), 2 (orange) and 4 (red) can be seen. On the right, log fold change in expression of “IRE1 induced” (*EEF2, TRPM8, SNHG8, VSTM2L*) and “IRE1 repressed” (*THBS1, LGI2, DDX11, HELLS, NEMP1*) genes when comparing increasing GG to each corresponding benign region. Each locally distinguishing comparison involves 331 GG1 spots, 938 GG2 spots, 1169 GG4 spots and 942, 703 and 1078 benign spots respectively. C) Correlation of expression pattern of the IRE1 activity gene set to radical prostatectomy Gleason score in the DKFZ Cancer Cell 2018 cohort. Statistical analysis was performed using Chi-squared test. Altered (red) status is defined as differentially significant z-score expression in tumour samples compared to the expression distribution of all log-transformed mRNA expression in adjacent normal samples in the cohort. D) Correlation of expression pattern of the IRE1 activity signature to pathology classification including adenocarcinoma (green), small cell (orange) and adenocarcinoma with NEPC features (purple). Statistical analysis was performed using Chi-squared test. Altered gene expression is defined as differentially significant log-transformed mRNA z-scores compared to mRNA signal from all samples in the cohort. Shortened versions of the IRE1 activity signature ((*VPS13A, EEF2, TRPM8, POLQ, TGM2, THBS1, LGI2, STX5, TRPC1*), (*VSTM2L, HELLS, CHI3L2, SNHG8, DDX11, SLITRK3, CRYL1, NEMP1, MYL9*)) tested for prognostication power showing significance in E) probability of overall survival and F) progression free survival in an mCRPC (PCF/SU2C) and primary disease (PRAD TCGA) cohort respectively. Altered expression defined as J) significant log-transformed mRNA z-scores compared to expression distribution of all samples (RNA Seq FPKM) and K) significant z-scores of tumour samples compared to the expression distribution of all log-transformed mRNA expression of adjacent normal samples in the cohort. Statistical analysis was performed using a Logrank (Mantel-Cox test). Four stars signify adjusted p value of <0.0001

Thereafter we correlated IRE1_18 expression with increasing grade group in this radical prostatectomy using GG1, GG2 and GG4 spots to recapitulate the full spectrum of multifocality and heterogeneity. The “IRE1 induced” component (*TRPM8, EEF2, SNHG8, VSTM2L*) increased whilst the “IRE1 repressed” component (*THBS1, DDX11, HELLS, NEMP1, LGI2*) decreased with tumour grade (Figure 6B).

To expand on this correlation, we explored data from 118 samples (DKFZ, Cancer Cell cohort) and observed significant differential expression changes in IRE1_18 genes with increasing Gleason score (Figure 6C). Furthermore, in the PCF/SU2C cohort, significant differential expression of these genes correlated with worsening pathology classification (NEPC/small cell disease) (Figure 6D). Indeed, change in mRNA status of these genes significantly correlated with clinical attributes in multiple datasets including NEPC scoring, BCR status and mutation count (Supplementary Table S25).

Since IRE1_18 represents different cell types and biological functions it should have specific prognostic power in different clinical settings. Indeed, splitting IRE1_18 in two subgroups comprising both “induced” and “repressed” genes significantly prognosticated both mCRPC (Figure 6E) and localised disease (Figure 6F) for overall and progression free survival respectively. In the PCF/SU2C cohort, patients with significantly differentially expressed mRNA of IRE1_18 had worse probability of overall survival compared to patients without (Supplementary Table S26). In the TCGA/PRAD cohort, patients with significantly differentially expressed mRNA of IRE1_18 showed significantly worse probability of progression free survival, compared to patients without (Supplementary Table S27).

Therefore, IRE1 activity represented by IRE1_18 showcases the importance of stress response biology in PCa pathophysiology, progression and treatment response.

## Discussion

We have previously reported that the IRE1-XBP1 axis of the unfolded protein response is regulated by androgen receptor activity and that inhibiting IRE1 restricts tumorigenesis in androgen-dependent models of prostate cancer. Sequencing well-annotated clinical samples has revealed that whilst AR activity is maintained in many cases as PCa progresses to metastasis, AR variants and target genes that support progression change. In addition, NEPC/DNPC PCa subtypes do not seem to rely on AR function or expression. By contrast, the UPR/IRE1 axis is capable of adapting as these disease states emerge. Given that the UPR-IRE1 axis is a feature of a wide range of cell types we have sought to determine how IRE1 activity relates to the emergence of acquired treatment resistance and to a spectrum of PCa sub-types or cell lineages. To assign IRE1 activity to cell lineages and PCa subtypes we derived a PCa-specific gene set (IRE1_18) by initially relying on data derived from PCa cell-lines and pre-clinical models. When refining IRE1_18 we selected genes that could discriminate between PCa sub-types in patient datasets and to prognosticate both localized and metastatic disease. De-convoluting IRE1_18 spatially, we determined that in clinical samples these genes are associated with a variety of cell types and biological functions likely to reflect niches in which treatment resistant disease can emerge.

Our cell-line/pre-clinical models cannot capture the cellular heterogeneity or biological complexity of these niches, but a potentially significant finding was that knocking out IRE1 in an androgen-dependent PCa cell-line results in the increased expression of lineage markers associated with club-like epithelial progenitor (*MMP7, PIGR, CP, LCN2*), and lineage reprogramming (*SOX2, EZH2, POU3F2*). IRE1 activity may impact cellular differentiation being associated with the acquisition of cancer stem cell characteristics in glioblastoma through XBP1s-driven repression of mir-148, which de-represses stemness TFs such as SOX2^38^. In PCa, *SOX2* has been associated with stemness and treatment resistance arising from knocking out TP53 and RB1 in androgen-dependent pre-clinical models^39^.

Intriguingly, knocking out IRE1 also leads to transcriptional changes resembling RB1 loss and oncogenic TF coregulation^40^ while correlating with AR loss and NEPC genotypes in mCRPC datasets. Importantly, pharmacologically increasing IRE1 activity in RB1 deficient cells sensitised them to physiological and pharmacological AR deprivation.

In the lifetime of PCa evolution and treatment, UPR/IRE1 activity has been shown to increase as an early stress-response, adaptive, pro-survival mechanism due to mounting metabolic stress and protein folding demand, parallel to the AR. But what happens when AR activity starts to decrease physiologically, or as a response to therapy? ADT-induced IRE1 signalling ablation may lead to adaptation that pushes a subset of tumours towards an AR+ CRPC or NEPC/DNPC phenotype, thus placing IRE1 activity at the centre of acquired treatment resistance.

Our study thus introduces two major novel themes for further investigation. Firstly, in more effectively stratifying patients for ADT and perhaps in the future, modulators of IRE1/UPR activity (ONC201^41^, NXP800^42^) by mapping IRE1 responses to AR antagonism (STAMPEDE), to metastatic site (Hartwig Medical Foundation) and to RB1-loss, BRCA2-amplification phenotypes lacking somatic RB1/BRCA2 changes (TCGA, SU2C). Secondly, for further developing models of lineage plasticity that can capture transition states between high AR activity, differentiated luminal epithelial cells and low AR activity, club-like epithelial progenitors aiming to predict the emergence of AR-negative basal or stem-like cells. IRE1 activity proxies (e.g., IRE1_18) representing multicellular tumoral niches are not dependent on accurate tumour sampling and can identify peri-tumoral tissue in false negative biopsies or inform risk stratification in histologically benign tissue with pre-tumoral genotypes. Indeed, clinically approved genomic tests (Decipher) contain genes attributed to multiple cell types and thus significantly associate with risk of metastasis and PCa specific mortality^43^.

The majority of significant work in this area of lineage plasticity has thus far focused on transcription factors (*ACSL1, POU3F2*) and chromatin-modifying enzymes (*EZH2*). Against this backdrop it will be interesting to determine whether changes in IRE1 activity create permissive states in which these more established drivers of lineage plasticity can function more effectively, or whether IRE1’s contribution is mechanistically independent.

## Summary

In conclusion, we explored PCa dynamic disease progression through the homeostatic mechanism of the UPR. By focusing on the UPR transducer IRE1, we produced an IRE1 activity gene set that can act as a proxy of AR activity, stress response, histological grade, cell type, clinical designation and potentially treatment response. IRE1 activity is commonly thought to be a driver of tumorigenesis. We here, report for the first time that adaptation to sustained stress and suppression of the IRE1 axis may induce lineage characteristics that mimic a club-like pre-cursor to RB1 loss and treatment resistant disease. This has major implications for IRE1 modulation in treatment resistant PCa as a differentiation therapy and for use of IRE1 activity as early identifier of lethal disease.

## Acknowledgments & Contributions

Dr Doultsinos and Prof. Mills designed the study, analysed data and wrote the manuscript. Dr Doultsinos carried out all experiments presented (qPCR, Western Blot, growth assays, viability assays, RNAseq) supported by Ms Abusamra, Ms Bridges and Prof. Hester. Ms Tomljanovic (bulk preclinical and clinical datasets), Dr Pilalis (bulk preclinical datasets), Dr Parmentier (single cell datasets), Dr Figiel (spatial datasets), Dr Zekri (ChIP-seq) and Dr Doultsinos (bulk clinical/preclinical, spatial datasets) carried out data analysis. Dr Leach and Prof. Bevan generated and provided PDEs. Prof. Chatziioannou, Dr Lamb, Dr Le Magnen and Prof. Zwart provided study design input and access to datasets and samples.

Special thanks to:

Dr Ursula Arndt for supporting sequencing experiments at the Granta Park Illumina Accelerator site. Dr Eric Chevet, Dr Tony Avril and Dr Diana Pelizarri (INSERM U1242) for providing IRE1 mutant constructs (Q780*) and MKC8866. Professor Nelson (Fred Hutchinson Cancer Research Centre) for providing LNCaP RB1 knockout and LNCaP RB1/TP53 knockout cell lines. Professor Zwart (NKI) for providing LNCaP ERN1 knockout cell lines. Dr Adam Sharp (ICR), Professor Scott Dehm (University of Minnesota), Dr Danielle Fairbrass (University of Oxford) and Dr Anette Magnussen (University of Oxford) for reading and commenting on the manuscript.

Author declaration: No generative AI was used in the preparation of this manuscript.

## Materials & Methods

### Cell lines & Culture Conditions

LNCaP and C4-2B cells were purchased from ATCC and cultured in RPMI supplemented with 10% foetal bovine serum (FBS) or DMEM supplemented with 10% foetal bovine serum (FBS) respectively. LNCaP knocked out for RB1 or knocked out for both RB1 and TP53 were kindly donated by Professor Peter Nelson (Fred Hutchinson Cancer Centre) and were cultured in RPMI supplemented with 10% foetal bovine serum (FBS). LNCaP knocked out for ERN1 were kindly donated by Professor Wilbert Zwart (The Netherlands Cancer Institute) and were cultured in RPMI supplemented with 10% foetal bovine serum (FBS). No antibiotics were used in the culturing of these cells as they may have the capacity to bind a variety of kinases including IRE1 as we previously reported^44^. For androgen deprived media, Charcoal Stripped FBS was used instead of normal FBS to supplement RPMI.

### Patient Derived Explant (PDE) tissue culture

PDEs were conducted with previously established protocols^11^, with changes to methodology for these studies. RPMI media supplemented with 5%FBS and 1mM enzalutamide or equivalent vehicle control (VC) was used to soak collagen dental sponges (Surgispon, AnserMedical). From consenting patients, tissue was collected from surgery and immediately transferred to the laboratory in ice-cold RPMI. Tissue was carefully dissected into 1mm3 cubes, and placed on to pre-soaked sponges in 24-well tissue culture plates. An additional 500ul of media with appropriate treatment was added to each well. After 72-hour culture, snapped frozen for RNA analysis. Additionally, matched pieces were formalin fixed and processed for histology as previously described^45^. RNA for RNA-seq was isolated using Monarch® kit according to manufacturer’s instructions (New England Biolabs).

Patient material was collected under Imperial College Healthcare Tissue Bank (REC 22/WA/0214) and the samples for this project (R18041) were issued from sub-collection reference number Uro_MW_13_010 from patients undergoing diagnosis of high-volume primary disease, with an age range between 55 and 88 years old.

### Primers and qPCR

RNA was extracted from cell pellets using TRIzol reagent (Fisher Scientific, #12034977) according to manufacturer’s instructions. cDNA was reverse transcribed from 2000ng of RNA (measured by Nanodrop) using Maxima Reverse Transcriptase (ThermoFisher Scientific EP0743) and associated reagents (RiboLock-Life TechnologIes-EO0381; DNTPs-Thermo Scientific-18427013; Random Hexamer-Life Technologies-SO142). qPCR was executed on a QuantStudio5 (Applied Biosystems) using Power SYBR Green PCR master mix (ThermoFisher Scientific) and results were analysed using the Design & Analysis Software 2.8.3 (Applied Biosystems).

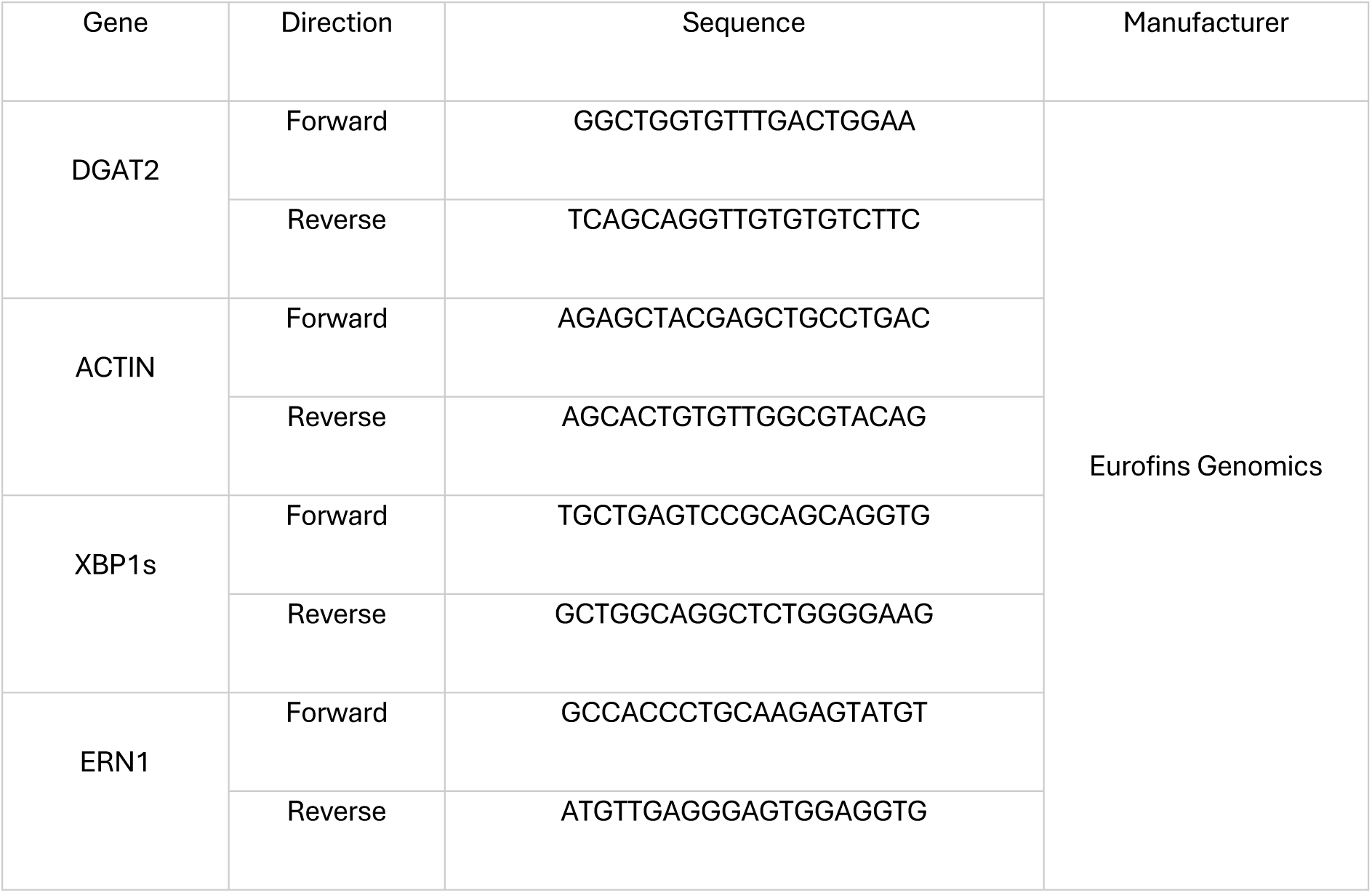
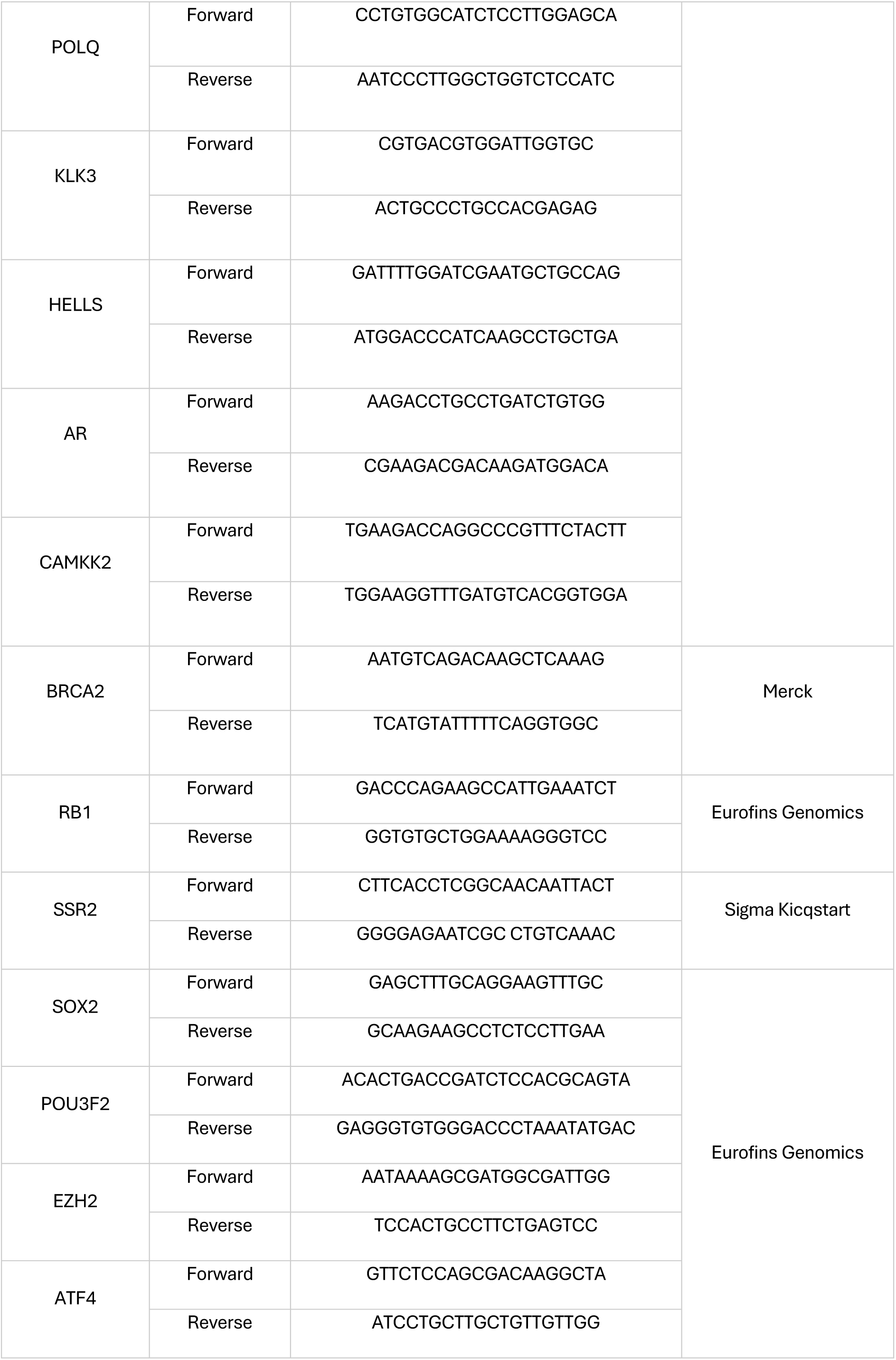
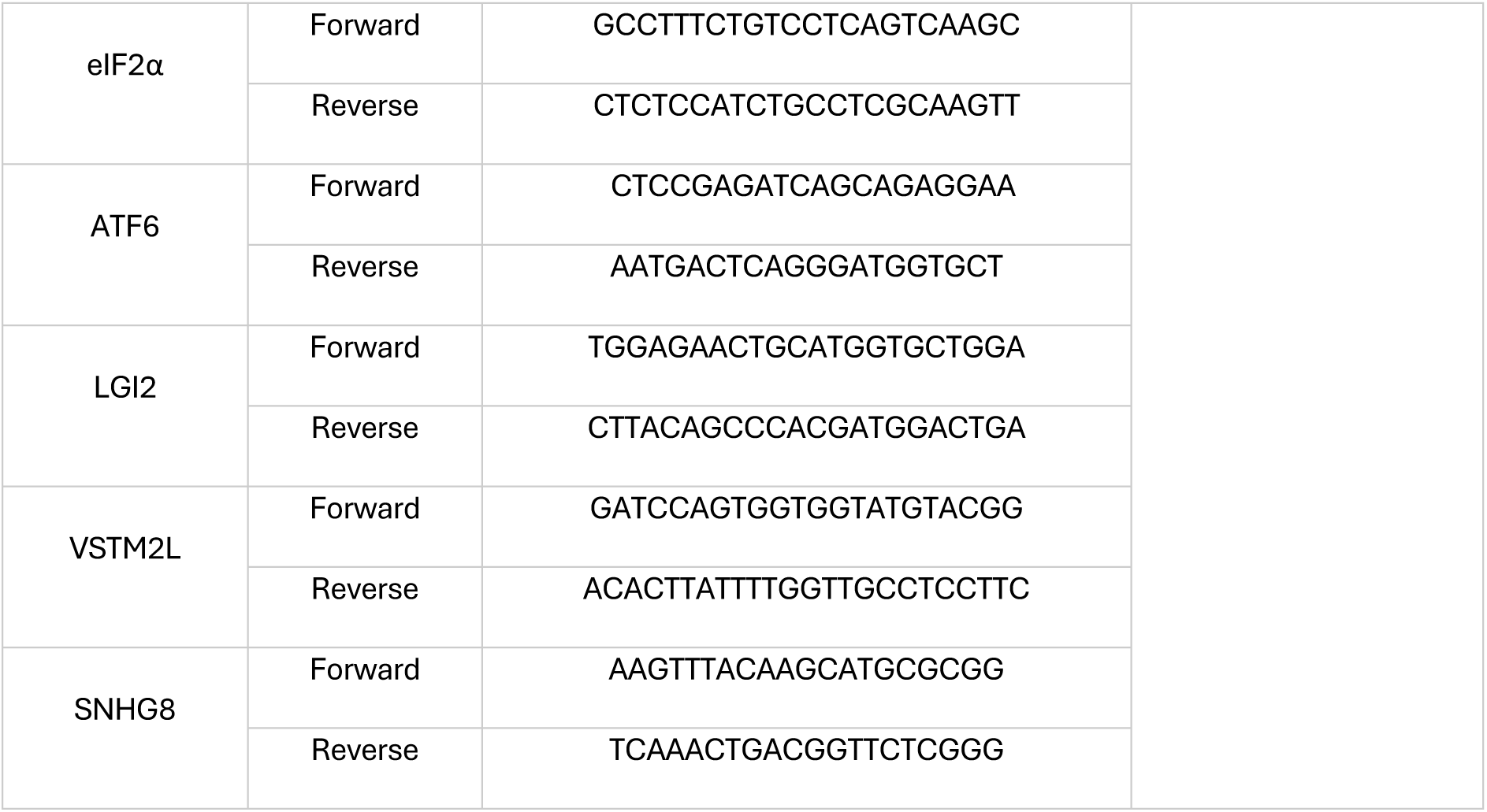

### Drug Treatments

Cells were seeded in 96, 24 or 6 well plate format depending on the assay (96 well plate for viability, 96 or 24 well plate for growth, 6 well plate for RNA extraction) and drugs were added using the range of concentrations below.

| Compound | Manufacturer | Lot number | Concentrations |
| --- | --- | --- | --- |
| IXA4 | Cayman Chemical | 0653659-8 | 10 - 25 μM |
| MKC8866 | Cayman Chemical | 0664887-3 | 5 μM |
| Thapsigargin | Sigma | 0000120608 | 100 nM |
| Tunicamycin | Sigma | 079M4004V | 500 nM |
| Enzalutamide | Selleck Chemicals | MDV3100 | 10 μM |

### Cell Titer Glo sensitivity assays

Cells were seeded into 96-well plates and incubated at 37 °C for 24 hours. They were then treated with drug combinations for a further 12 hours. Percentage survival was measured photometrically using the CellTiter-Glo 2.0 viability assay (Promega) in a microplate reader (FLUOstar OMEGA 415-2812). Survival was normalized to the control treatment.

### Growth assays

Cells were seeded into 96-well plates and incubated at 37 °C for 24 hours. Complete media change followed to either full (FM) or charcoal stripped media (CSM) for 120 hours. Cell growth was monitored using Incucyte technology taking confluence measurements every 24 hours. Growth was normalised to the initial measurement at time 0.

### Transfection assays and RNA silencing

For stable expression of IRE1 mutant Q780*, LNCaP ERN1^-/-^ cells were transfected with the Q780* construct^14^ using LTX Plus reagent (ThermoFisher Scientific, 15338100) according to the manufacturer’s instructions. Successfully transfected cells were selected by cell sorting for GFP expression using a BD FACSAriaIII cell sorter. Thereafter, constant GFP expression was monitored using the ZOE^TM^ Fluorescent Cell Imager. Lipofectamine RNAiMax transfection reagent (Thermofisher Scientific, 13778150) was used for transfecting siRNAs. Cells were incubated at 37 °C for 48 hours before extracting RNA for downstream analysis.

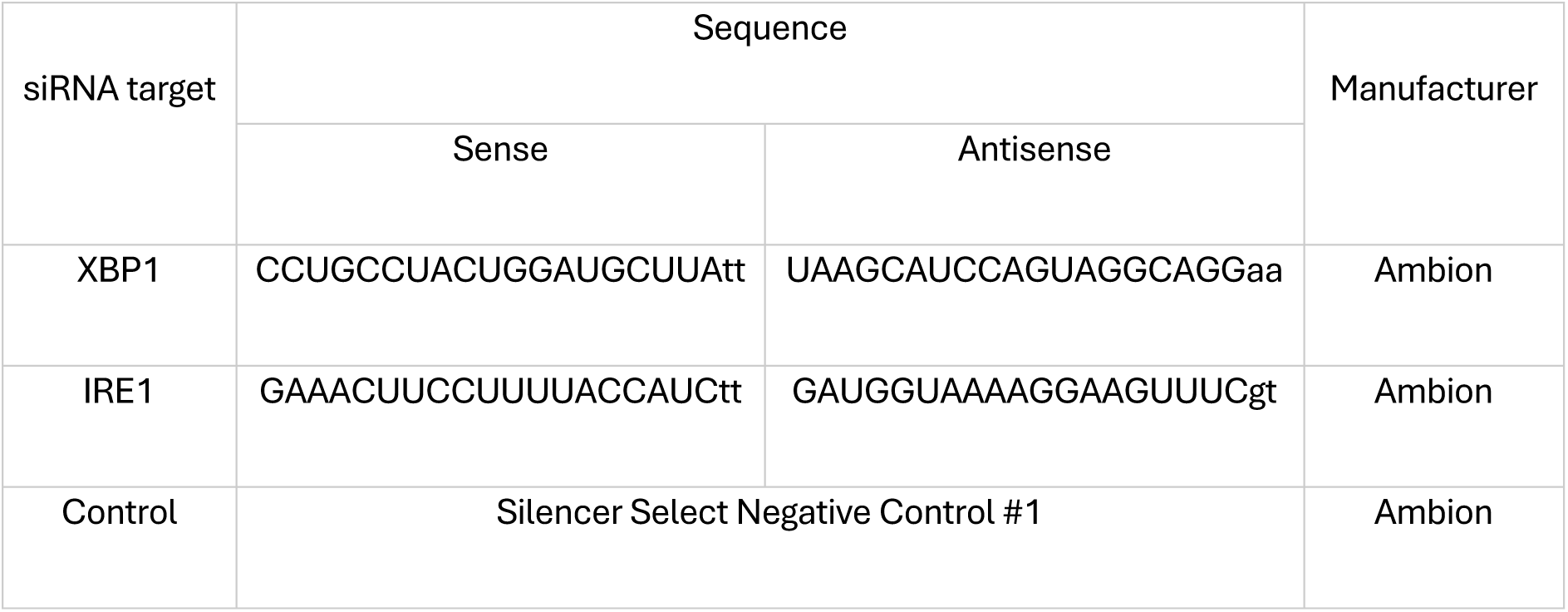

### Immunoblotting and Antibodies

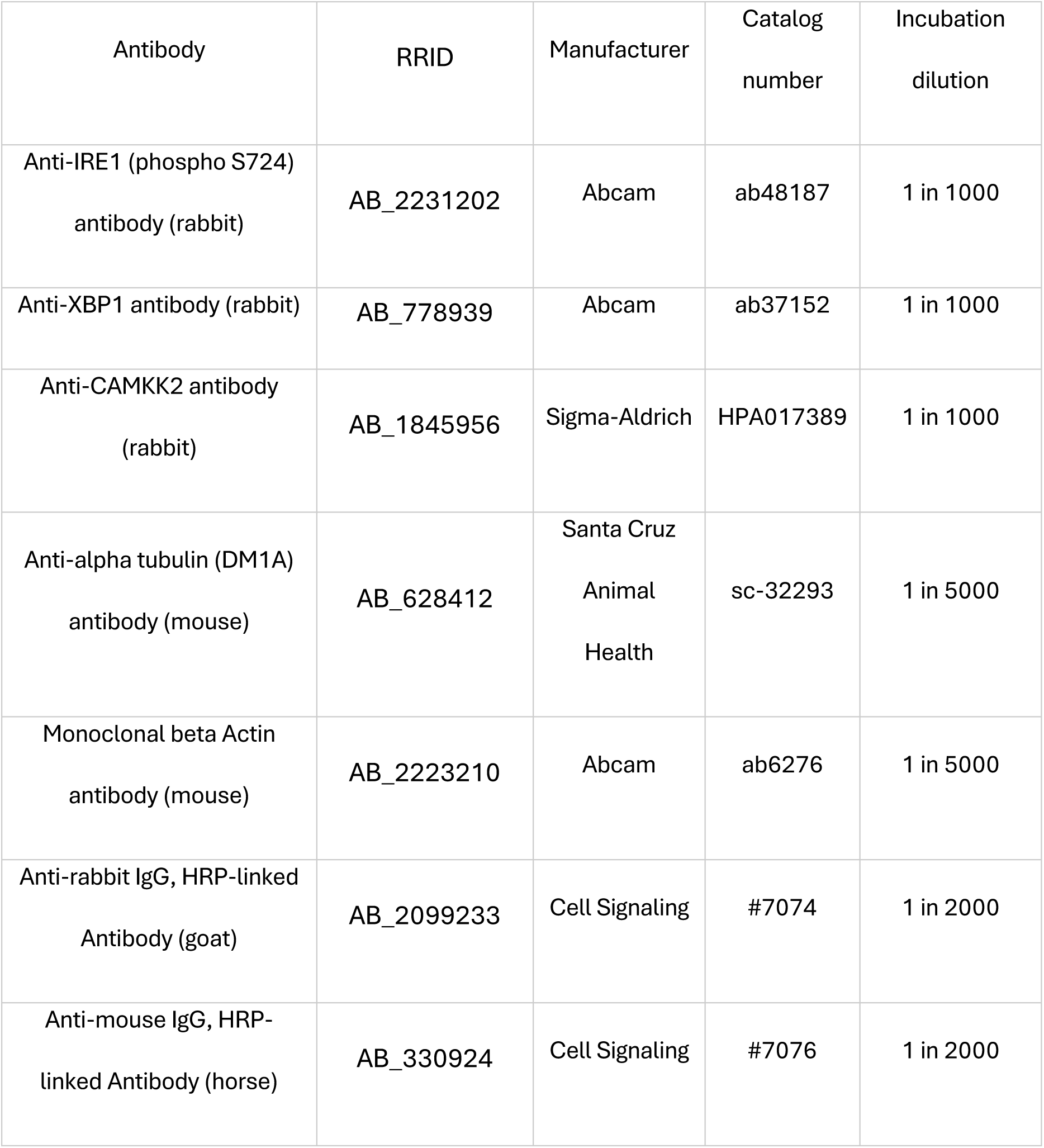

| Antibody | RRID | Manufacturer | Catalog number | Incubation dilution |
| --- | --- | --- | --- | --- |
| Anti-IRE1 (phospho S724) antibody (rabbit) | AB_2231202 | Abcam | ab48187 | 1 in 1000 |
| Anti-XBP1 antibody (rabbit) | AB_778939 | Abcam | ab37152 | 1 in 1000 |
| Anti-CAMKK2 antibody (rabbit) | AB_1845956 | Sigma-Aldrich | HPA017389 | 1 in 1000 |
| Anti-alpha tubulin (DM1A) antibody (mouse) | AB_628412 | Santa Cruz<br>Animal Health | sc-32293 | 1 in 5000 |
| Monoclonal beta Actin antibody (mouse) | AB_2223210 | Abcam | ab6276 | 1 in 5000 |
| Anti-rabbit IgG, HRP-linked Antibody (goat) | AB_2099233 | Cell Signaling | #7074 | 1 in 2000 |
| Anti-mouse IgG, HRP-linked Antibody (horse) | AB_330924 | Cell Signaling | #7076 | 1 in 2000 |

### Online Tools/Databases

- <u>cBioportal (RRID:SCR_014555)</u>: RNA sequencing datasets investigated included the Metastatic Prostate Adenocarcinoma (SU2C/PCF Dream Team, PNAS 2019^46^), Prostate Cancer (DKFZ, Cancer Cell 2018^47^), Prostate Adenocarcinoma (MSK, Cancer Cell 2010^48^) and Prostate Adenocarcinoma (TCGA, Firehose Legacy or PanCancer). The comparisons were based on log-transformed mRNA expression z-scores compared to expression distribution of all samples (RNA Seq FPKM) or of tumour samples compared to the expression distribution of all adjacent normal samples in the cohort when available with a z score threshold of +/- 2.0. Altered mRNA was defined as mRNA significantly high or low compared to all or normal samples in each dataset using the z score threshold. Kaplan Meier curves were generated for the Prostate Adenocarcinoma (SU2C/PCF Dream Team, PNAS 2019) and Prostate Adenocarcinoma (TCGA SCR_003193, PanCancer) datasets. Only samples from patients with survival data or progression data were used in the generation of these curves respectively. Samples with any significant alteration in mRNA expression were compared to samples with zero significant changes in the former whilst samples with significant alterations in at least three genes were compared to samples with zero significant changes in the latter.
- <u>Prostate Cancer Atlas</u>^32^: DESeq2 differential expression analysis was carried out foreach of sample groups designated as Primary, AR+ CRPC, NEPC and DNPC compared to the sample group designated as normal tissue. Differentially significant mRNA expression changes (padj</=0.01) were documented to mine for genes of interest representing IRE1 activity. Pseudotime disease progression plots were generated for “IRE1 induced” and “IRE1 repressed” sub-groups of the IRE1 activity gene set using the in-built functionality of the Prostate Cancer Atlas (https://prostatecanceratlas.org/app/home). Further in-built functionality was used to analyse gene set enrichment of our IRE1 activity gene set in the normal, primary, AR+ CRPC, NEPC and DNPC samples groups.
- <u>Correlation AnalyzeR</u>^49^: The online interface of this tool was used to verify the correlation of the IRE1 activity gene set with other significantly differentially expressed genes in prostate cancer and the biological processes it represents.
- <u>CAMCAPP</u>^50^: This tool was used to verify gene by gene correlation of the IRE1 activity gene set with clinical co-variates in three separate clinical cohorts (Stockholm, Cambridge, MSKCC).

### RNA extraction, library prep and sequencing

RNA was extracted from cell line pellets using the RNeasy Plus Mini kit. Samples were QC’d using Nanodrop and the Qubit Flex High Sensitivity ID kit. Libraries were prepared using the TruSeq Stranded Total RNA with Illumina Ribo-Zero plus rRNA Depletion workflow (https://emea.support.illumina.com/downloads/truseq-stranded-total-rna-with-ribo-zero-plus-rrna-depletion-ref-guide.html). Index adapters, rRNA depletion kit were acquired in addition to the TruSeq Stranded Total Library Prep Gold kit (https://emea.illumina.com/products/by-type/sequencing-kits/library-prep-kits/truseq-stranded-total-rna.html#tabs-bb5d9c771f-item-01fd0b3d08-order). cDNA library QC was carried out on Tapestation and Libraries were pooled, diluted to ∼10 nM, denatured, and further diluted for paired-end sequencing (NovaSeq6000 platform, Illumina, NovaSeq 6000 S2/S4 reagent kit v1.5, 300 cycles). A test run prior to full sequencing was executed on a MiSeq system. Bulk paired-end RNA sequencing was performed to obtain at least 20mn reads per sample. Raw data were processed using a standardized computational workflow to obtain gene-level raw counts. FASTQ files were quality-checked using FastQC to assess read quality and adapter contamination. High-quality reads were aligned to the human reference genome (GRCh38, Ensembl release 108) using STAR aligner (Dobin et al., 2013). Gene-level quantification was performed using HTSeq-count (Anders et al., 2015).

### RNA Sequencing Data analysis

1. Spatial transcriptomics data were obtained from a previously published dataset derived from radical prostatectomy tissue of a patient diagnosed with multifocal prostate cancer (Nature). Each spatial transcriptomics spot was reviewed by two independent pathologists as part of a consensus approach for pathological annotation.

- <u>Virtual Biopsy</u>: The analysis was performed using the same approach as previously described (Molecular Cancer). Gene expression data were extracted from the barcode ID of designed biopsies (LoupeBrowser, v8.0.0) using R (v4.4.0). Libraries were normalised using the spaceranger aggr (v2.0) function, which corrects for batch effects to facilitate the comparison of gene expression data across samples and minimise technical variability. The analysis included seven distinct tissue sections, representing a total of 3721 spatial transcriptomics spots. To maintain data integrity, spots with fewer than 500 Unique Molecular Identifier (UMI) counts were removed. Fold changes were calculated using EdgeR (v3.40.1). as previously described^37^. <u>Single Visium 6.5mm squares</u>: The raw sequencing reads were directly processed using Space Ranger and mapped using reference genome GRCh38, release 93 as previously described^34^. Each histological square was treated as a distinct spatial transcriptomic experiment to determine if, in an independent manner, gene sets could be used as distinguishing tools between histological characteristics. Histological annotations (benign, GG1, GG2, GG4, stroma) were compared using either the locally distinguishing or globally distinguishing functions of the Loupe Browser which was also used to generate ST images displayed.
2. Single Cell PCa tissue data from Chen et al. were accessed via the GEO online portal (GSM4203181) as a raw count matrix. Quality control filtering was applied based on the criteria described in the original publication: cells with fewer than 200 or more than 5,218 detected genes, more than 20% mitochondrial content, and fewer than 56 housekeeping genes were discarded. The filtered data were then converted into an SCE object, separated by patient ID, and normalized using a cluster-based normalization factor computed after each sample was first partitioned into biologically relevant clusters using the quickCluster(min.size = 100) function from the scran R package (v1.26.2). Next, to correct for differences in sequencing depth across batches, all SCE objects were rescaled using the MultiBatchNorm(min.mean = 0.1) function from the Batchelor R package (v1.14.1). Epithelial cells were retrieved using the singleR package (v2.0.0), leveraging the BlueprintEncodeData reference accessed via the celldex package (v1.16.0). Using scran (v1.26.2), variance modeling of gene expression was performed for each sample individually with the modelGeneVar() function; results were merged with combineVar(), and the top 1,000 most highly variable genes were selected using getTopHVGs() for subsequent analysis. Batch correction was applied using the fastMNN(k = 300, d = 20) function from the Batchelor package. To separate luminal malignant cells from healthy epithelial cell types, clustering was performed on the integrated data using the clusterCells() function from the scran R package (v1.26.2). The clustering was conducted with the Walktrap algorithm, exploring different values of k = 10, which resulted in 14 distinct clusters. All clusters were considered tumor except cluster 7 and 14, which were negative of both tumor and luminal markers (e.g AMACR, PCA3, KLK3) and were positive for basal markers such as KRT19, KRT18 and KRT14. PCa tissue data from Hirz et al. were accessed via the GEO online portal (GSE181294) as a raw count matrix. Cells passing quality thresholds and cell type annotation, as defined in the original publication, were retrieved from the annotation table provided by the authors and stored in an SCE object. Similar to the Chen et al. dataset, each sample first underwent cluster-based normalization followed by inter-sample correction to account for sequencing depth differences between batches.
3. Bulk The RNA sequencing data (raw counts) of the Labrecque mCRPC patient cohort (accession number GSE126078) and the RB1^-/-^/TP53^-/-^ cell line dataset (accession number GSE147250) was retrieved from GEO. TCGA data was accessed via the GDC Data Portal (https://portal.gdc.cancer.gov/projects/TCGA-PRAD, TCGA-PRAD.2022-04-27). The Beltran patient cohort FPKM data was obtained from the supplementary material of Beltran et al., 2016. Data visualization and statistical analyses of the bulk RNA sequencing datasets were performed using R statistical software (v4.4.1). Batch effect correction was executed using the sva package (v.3.54.0). Differential gene expression analysis was performed using the DESeq2 package (v1.40.2), with subsequent LFC shrinkage using the ‘apeglm’ estimator. Gene set enrichment analysis (GSEA) was conducted on a subset of the Molecular Signatures Database (MSigDB) v2023.2.Hs using the clusterProfiler package (v4.8.2). Gene Set Variation Analysis (GSVA) was used for single-sample pathway enrichment as implemented in the GSVA package (v1.48.3). Differential expression at the pathway level was assessed using the lmFit function from limma (v3.62.2). For visualization and clustering purposes, the variance stabilizing transformation (VST) was applied using the DESeq2 package. Visualization was performed using ggplot2 (v3.5.0) and heatmaps were generated using the pheatmap package (v1.0.12). The viper R package (v1.4.0) was used for single-sample regulon enrichment and differential testing of regulon activity levels between sample groups. The regulons of 1695 human transcription factors were previously inferred using ARACNE -AP on an in-house set of mCRPC patient tumors. Unless otherwise specified, the statistical significance threshold across all analyses was set at 0.05 and a Benjamini–Hochberg correction was applied to account for multiple testing. The data discussed in this publication have been deposited in NCBI’s Gene Expression Omnibus (Edgar et al., 2002) and are accessible through GEO Series accession number GSE297633 (https://www.ncbi.nlm.nih.gov/geo/query/acc.cgi?acc=GSE297633).

## References

1. Doultsinos, D. & Mills, I. The role of the androgen receptor as a driver and mitigator of cellular stress. J Mol Endocrinol 1, (2020).

2. Rawla, P. Epidemiology of Prostate Cancer. World J Oncol 10, 63–89 (2019).

3. Almanza, A., et al. Endoplasmic reticulum stress signalling – from basic mechanisms to clinical applications. FEBS Journal vol. 286 241–278 Preprint at 10.1111/febs.14608 (2019).

4. Almanza, A. et al. Regulated IRE1α-dependent decay (RIDD)-mediated reprograming of lipid metabolism in cancer. Nature Communications 2022 13:1 13, 1–13 (2022).

5. Bright, M. D., Itzhak, D. N., Wardell, C. P., Morgan, G. J. & Davies, F. E. Cleavage of BLOC1S1 mRNA by IRE1 Is Sequence Specific, Temporally Separate from XBP1 Splicing, and Dispensable for Cell Viability under Acute Endoplasmic Reticulum Stress. Mol Cell Biol 35, 2186–2202 (2015).

6. Raymundo, D. P. et al. Pharmacological Targeting of IRE1 in Cancer. Trends Cancer 6, 1018–1030 (2020).

7. Sharma, N. L. et al. The Androgen Receptor Induces a Distinct Transcriptional Program in Castration-Resistant Prostate Cancer in Man. Cancer Cell 23, 35–47 (2013).

8. Sheng, X. et al. IRE1α-XBP1s pathway promotes prostate cancer by activating c-MYC signaling. Nat Commun 10, 323 (2019).

9. Sheng, X. et al. Divergent androgen regulation of unfolded protein response pathways drives prostate cancer. EMBO Mol Med 7, 788–801 (2015).

10. Hamdy, F. C. et al. Fifteen-Year Outcomes after Monitoring, Surgery, or Radiotherapy for Prostate Cancer. New England Journal of Medicine 388, 1547–1558 (2023).

11. Centenera, M. M. et al. A patient-derived explant (PDE) model of hormone-dependent cancer. Mol Oncol 12, 1608 (2018).

12. Conteduca, V. et al. Clinical features of neuroendocrine prostate cancer. Eur J Cancer 121, 7–18 (2019).

13. Stelloo, S. et al. Androgen modulation of XBP1 is functionally driving part of the AR transcriptional program. Endocr Relat Cancer 27, 67–79 (2020).

14. Lhomond, S. et al. Dual IRE1 RNase functions dictate glioblastoma development. EMBO Mol Med 10, (2018).

15. Mu, P. et al. SOX2 promotes lineage plasticity and antiandrogen resistance in TP53-and RB1-deficient prostate cancer. Science (1979) 355, (2017).

16. Venkadakrishnan, V. B. et al. Lineage-specific canonical and non-canonical activity of EZH2 in advanced prostate cancer subtypes. Nature Communications 2024 15:1 15, 1–15 (2024).

17. Miao, C. et al. RB1 loss overrides PARP inhibitor sensitivity driven by RNASEH2B loss in prostate cancer. Sci Adv 8, 9794 (2022).

18. Flores, M. & Goodrich, D. W. Retinoblastoma Protein Paralogs and Tumor Suppression. Front Genet 13, 818719 (2022).

19. Hirz, T. et al. Dissecting the immune suppressive human prostate tumor microenvironment via integrated single-cell and spatial transcriptomic analyses. Nature Communications 2023 14:1 14, 1–20 (2023).

20. Kiviaho, A. et al. Single cell and spatial transcriptomics highlight the interaction of club-like cells with immunosuppressive myeloid cells in prostate cancer. Nature Communications 2024 15:1 15, 1–16 (2024).

21. Huang, F. W. et al. Club-like cells in proliferative inflammatory atrophy of the prostate. J Pathol 261, 85–95 (2023).

22. Song, H. et al. Single-cell analysis of human primary prostate cancer reveals the heterogeneity of tumor-associated epithelial cell states. Nature Communications 2022 13:1 13, 1–20 (2022).

23. Joseph, D. B. et al. 5-Alpha reductase inhibitors induce a prostate luminal to club cell transition in human benign prostatic hyperplasia. J Pathol 256, 427–441 (2022).

24. Ku, S. Y. et al. Rb1 and Trp53 cooperate to suppress prostate cancer lineage plasticity, metastasis, and antiandrogen resistance. Science (1979) 355, (2017).

25. Mu, P. et al. SOX2 promotes lineage plasticity and antiandrogen resistance in TP53-and RB1-deficient prostate cancer. Science (1979) 355, 84–88 (2017).

26. Nyquist, M. D. et al. Combined TP53 and RB1 Loss Promotes Prostate Cancer Resistance to a Spectrum of Therapeutics and Confers Vulnerability to Replication Stress. Cell Rep 31, (2020).

27. Logue, S. E. et al. Inhibition of IRE1 RNase activity modulates the tumor cell secretome and enhances response to chemotherapy. Nat Commun 9, 3267 (2018).

28. Grandjean, J. M. D. et al. Pharmacologic IRE1/XBP1s activation confers targeted ER proteostasis reprogramming. Nature Chemical Biology 2020 16:10 16, 1052–1061 (2020).

29. Labrecque, M. P. et al. Molecular profiling stratifies diverse phenotypes of treatment-refractory metastatic castration-resistant prostate cancer. J Clin Invest 129, 4492 (2019).

30. Thangavel, C. et al. RB loss promotes prostate cancer metastasis. Cancer Res 77, 982– 995 (2017).

31. He, M. X. et al. Transcriptional mediators of treatment resistance in lethal prostate cancer. Nature Medicine 2021 27:3 27, 426–433 (2021).

32. Bolis, M. et al. Dynamic prostate cancer transcriptome analysis delineates the trajectory to disease progression. Nature Communications 2021 12:1 12, 1–15 (2021).

33. Aggarwal, R. et al. Prognosis Associated With Luminal and Basal Subtypes of Metastatic Prostate Cancer Supplemental content. JAMA Oncol 7, 1644–1652 (2021).

34. Erickson, A. et al. Spatially resolved clonal copy number alterations in benign and malignant tissue. Nature 2022 608:7922 608, 360–367 (2022).

35. Chen, S. et al. Single-cell analysis reveals transcriptomic remodellings in distinct cell types that contribute to human prostate cancer progression. Nat Cell Biol 23, 87–98 (2021).

36. Bluemn, E. G. et al. Androgen Receptor Pathway-Independent Prostate Cancer Is Sustained through FGF Signaling. Cancer Cell 32, 474 (2017).

37. Figiel, S. et al. Spatial transcriptomic analysis of virtual prostate biopsy reveals confounding effect of tissue heterogeneity on genomic signatures. Mol Cancer 22, 1–5 (2023).

38. Doultsinos, D., et al. Constitutive UPRER activation sustains tumor cell differentiation. bioRxiv (2019) doi:10.1101/594630.

39. Venkadakrishnan, V. B., Yamada, Y., Weng, K., Idahor, O. & Beltran, H. Significance of RB loss in unlocking phenotypic plasticity in advanced cancers. Mol Cancer Res 21, 497 (2023).

40. Mandigo, A. C. et al. Novel Oncogenic Transcription Factor Cooperation in RB-Deficient Cancer. Cancer Res 82, 221–234 (2022).

41. Amoroso, F. et al. Modulating the unfolded protein response with ONC201 to impact on radiation response in prostate cancer cells. Scientific Reports 2021 11:1 11, 1–16 (2021).

42. Welti, J. et al. NXP800 activates the unfolded protein response, altering AR and E2F function to impact castration-resistant prostate cancer growth. Clin Cancer Res OF1–OF18 (2025) doi:10.1158/1078-0432.CCR-24-2386/750830/AM/NXP800-ACTIVATESTHE-UNFOLDED-PROTEIN-RESPONSE.

43. Feng, F. Y. et al. Validation of a 22-Gene Genomic Classifier in Patients With Recurrent Prostate Cancer: An Ancillary Study of the NRG/RTOG 9601 Randomized Clinical Trial. JAMA Oncol 7, 544 (2021).

44. Doultsinos, D. et al. Peptidomimetic-based identification of FDA-approved compounds inhibiting IRE1 activity. FEBS Journal 288, 945–960 (2021).

45. Leach, D. A. et al. Simultaneous inhibition of TRIM24 and TRIM28 sensitises prostate cancer cells to antiandrogen therapy, decreasing VEGF signalling and angiogenesis. Mol Oncol (2025) doi:10.1002/1878-0261.70065.

46. Abida, W. et al. Genomic correlates of clinical outcome in advanced prostate cancer. Proc Natl Acad Sci U S A 166, 11428–11436 (2019).

47. Gerhauser, C. et al. Molecular Evolution of Early-Onset Prostate Cancer Identifies Molecular Risk Markers and Clinical Trajectories. Cancer Cell 34, 996–1011.e8 (2018).

48. Taylor, B. S. et al. Integrative genomic profiling of human prostate cancer. Cancer Cell 18, 11–22 (2010).

49. Miller, H. E. & Bishop, A. J. R. Correlation AnalyzeR: functional predictions from gene co-expression correlations. BMC Bioinformatics 22, (2021).

50. Dunning, M. J. et al. Mining Human Prostate Cancer Datasets: The “camcAPP” Shiny App. EBioMedicine 17, 5 (2017).

