## Supplementary Figures for "IRE1 activity regulates tumour and microenvironment cell lineage states while stratifying localised and metastatic prostate cancer"

Supplementary figure S1

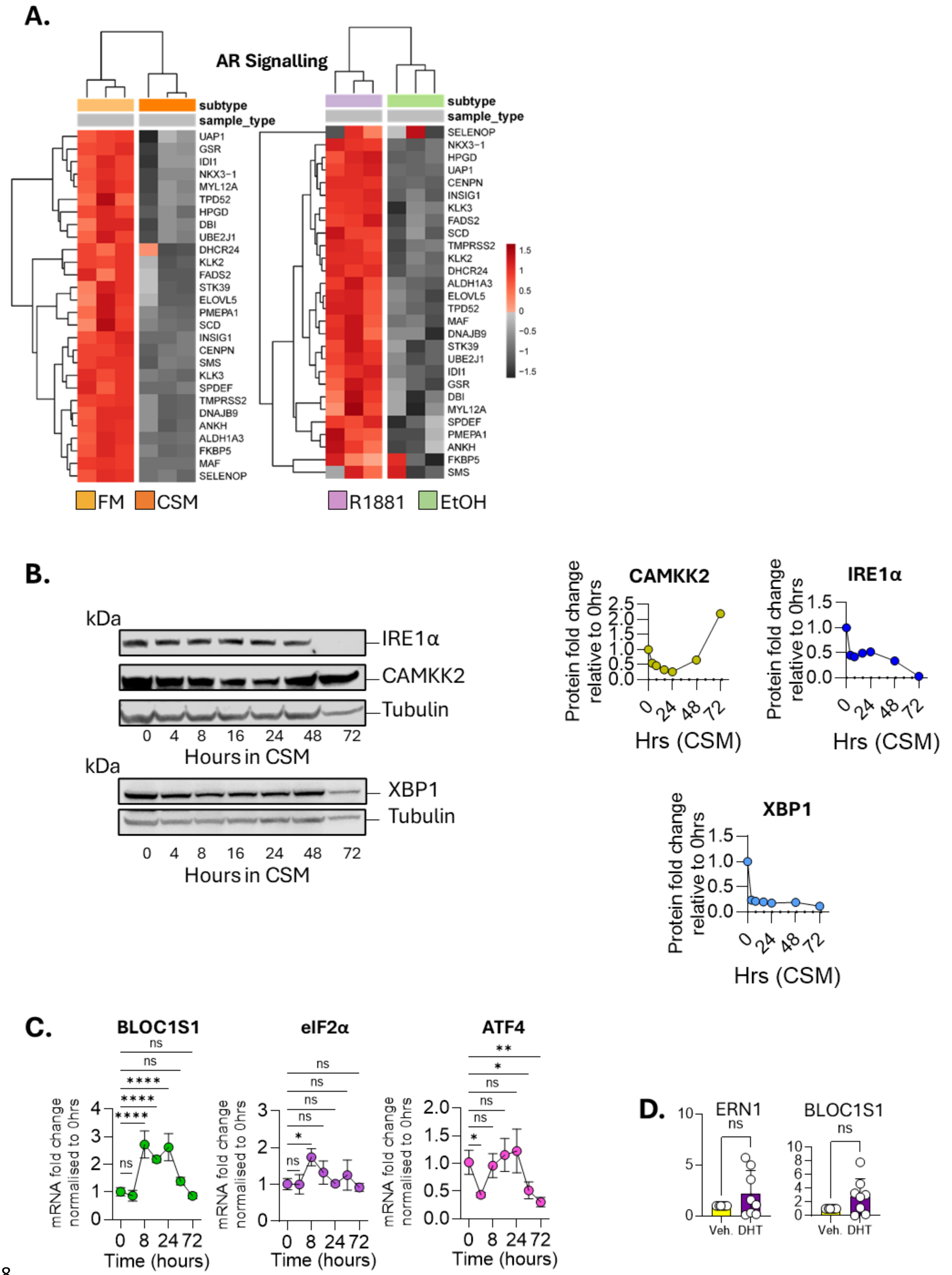

**Supplementary Figure S1**. A) Heatmaps showing consensus AR signalling genes upregulated in the R1881 (purple) condition compared to EtOH (green) and downregulated in the CSM (orange) condition compared to FM (yellow). B) Blots of lysates derived from LNCaP cells grown in CSM for 72 hours showing protein levels of IRE1α, CAMKK2, XBP1 and Tubulin with accompanying densitometry. Each time point was normalised to the housekeeping tubulin and then the values normalised to the 0-hour time point for each of IRE1, XBP1 and CAMKK2. C) mRNA fold change of IRE1-RIDD (*BLOC1S1*) and PERK activity (*eIF2α, ATF4*) genes in LNCaP parental cells grown over 72 hours in CSM normalised to FM control. Statistical analysis was performed using an ordinary one-way ANOVA with Dunnett's multiple comparisons test and single pooled variance. D) mRNA fold change of BLOC1S1, ERN1 in patient derived explants (PDEs) treated with dihydrotestosterone (DHT) normalised to PDEs treated with vehicle control. Statistical analysis was performed using a T-test per condition. Each experiment was performed at least in duplicate. Four stars signify adjusted p value of <0.0001.

#### Supplementary figure S2

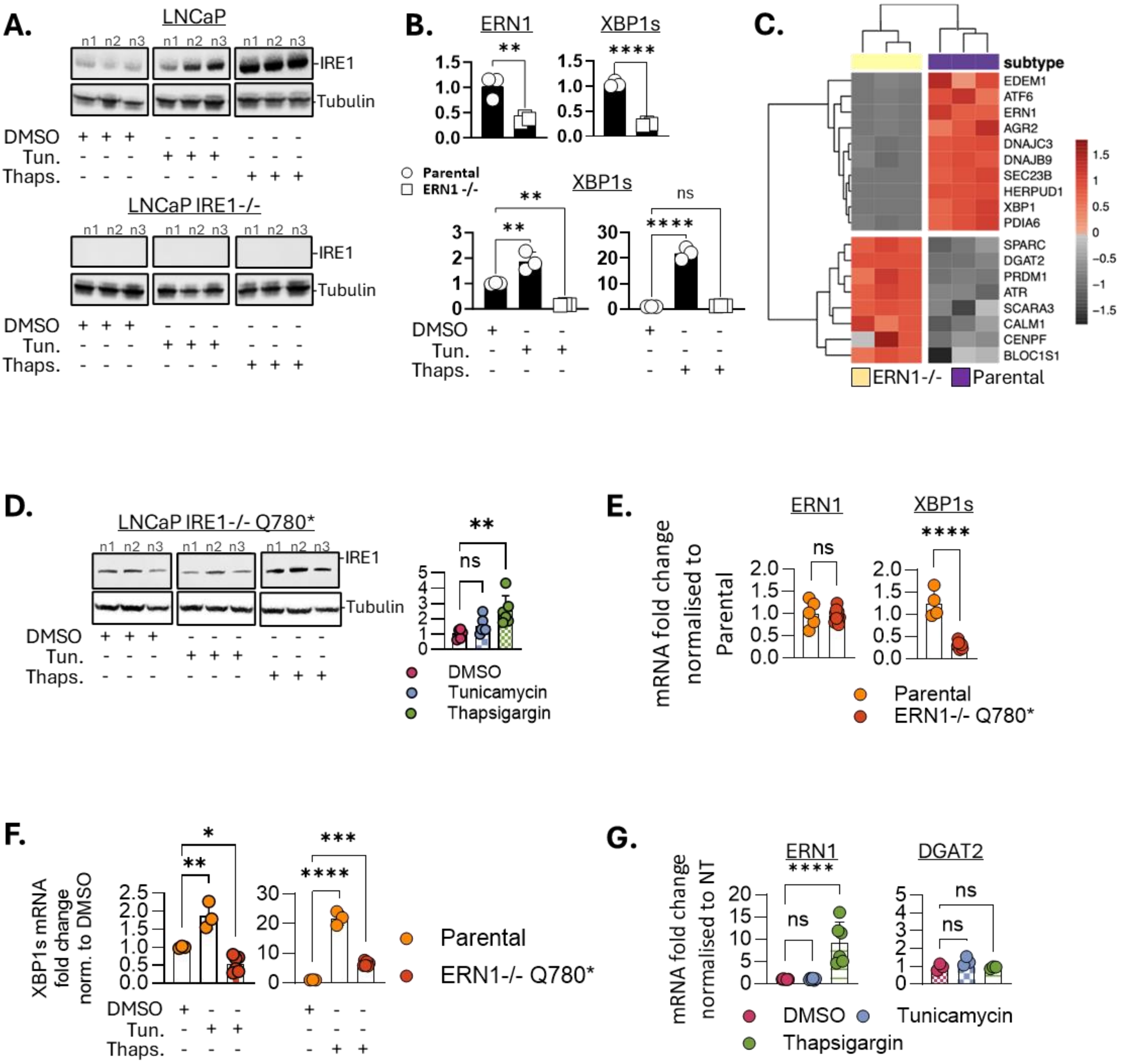

**Supplementary Figure S2**. A) Western blots in triplicate for IRE1α of LNCaP parental (top) or LNCaP ERN1$^{-/-}$ cells (bottom) in the presence of ER stressors tunicamycin or thapsigargin (100nM). The DMSO control is also shown. B) mRNA fold change of ERN1, XBP1s in basal conditions (FM, top) or in the presence of tunicamycin or thapsigargin (bottom) in LNCaP parental or LNCaP ERN1$^{-/-}$ cells. Top: Statistical analysis was performed using T-tests. Bottom: Statistical analysis was performed using an ordinary one-way ANOVA with Dunnett's multiple comparisons test and single pooled variance. C) Heatmap of canonical IRE1-XBP1 or IRE1-RIDD targets in RNAseq data derived from LNCaP parental (purple) or LNCaP ERN1-/- (yellow) cells. D) Western blots in triplicate for IRE1α of LNCaP ERN1-/- Q780* cells in the presence of ER stressors tunicamycin or thapsigargin (100nM). The DMSO control is also shown. Accompanying densitometry measures the impact of ER stress on protein levels of IRE1 mutant protein. E) mRNA fold change of ERN1 and XBP1s in LNCaP parental and LNCaP ERN1$^{-/-}$ Q780* cells under basal conditions (FM). Statistical analysis was performed using a T-test per condition. F) IRE1 induced activity marker XBP1s mRNA fold change in LNCaP parental or LNCaP ERN1$^{-/-}$ Q780* cells treated with DMSO, Tunicamycin or Thapsigargin normalised to parental cells treated with DMSO. Statistical analysis was performed using an ordinary one-way ANOVA with Dunnett's multiple comparisons test and single pooled variance. G) IRE1 repressed activity marker *DGAT2* and *ERN1* mRNA fold change in LNCaP ERN1$^{-/-}$ Q780* cells treated with DMSO, Tunicamycin or Thapsigargin normalised the DMSO condition. Statistical analysis was performed using an ordinary one-way ANOVA with Dunnett's multiple comparisons test and single pooled variance. Each experiment was performed at least in duplicate. Four stars signify adjusted p value of <0.0001.

### Supplementary figure S3

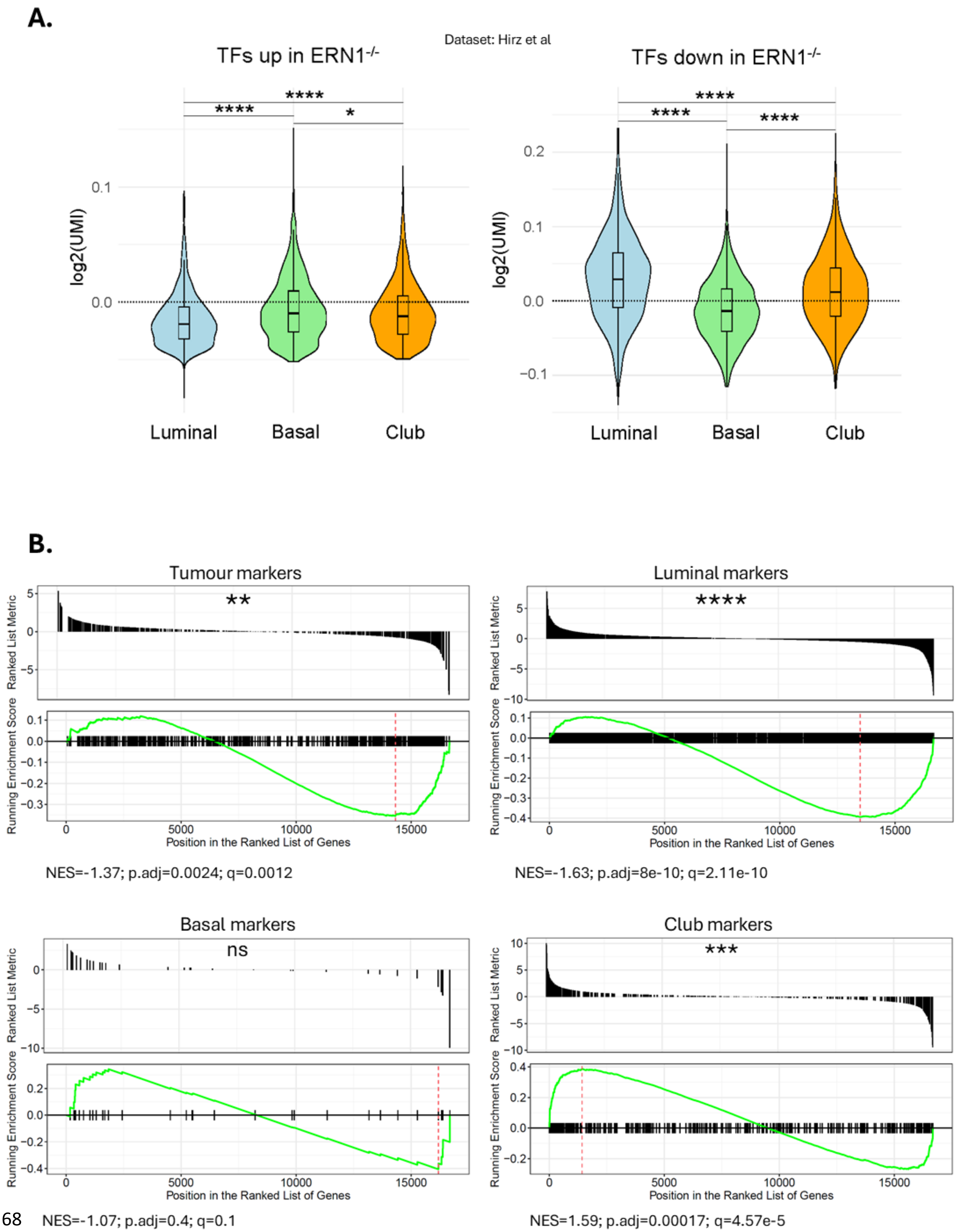

**Supplementary Figure S3**. A) Violin plots showing the enrichment of transcription factors significantly upregulated or downregulated in ERN1$^{-/-}$ cells, in basal cells and club cells compared to luminal epithelial cells in a single cell dataset. B) Gene set enrichment plots of gene sets representing tumour, luminal epithelial, basal epithelial and club epithelial cells in the LNCaP ERN1$^{-/-}$ cells compared to parental.

#### Supplementary figure S4

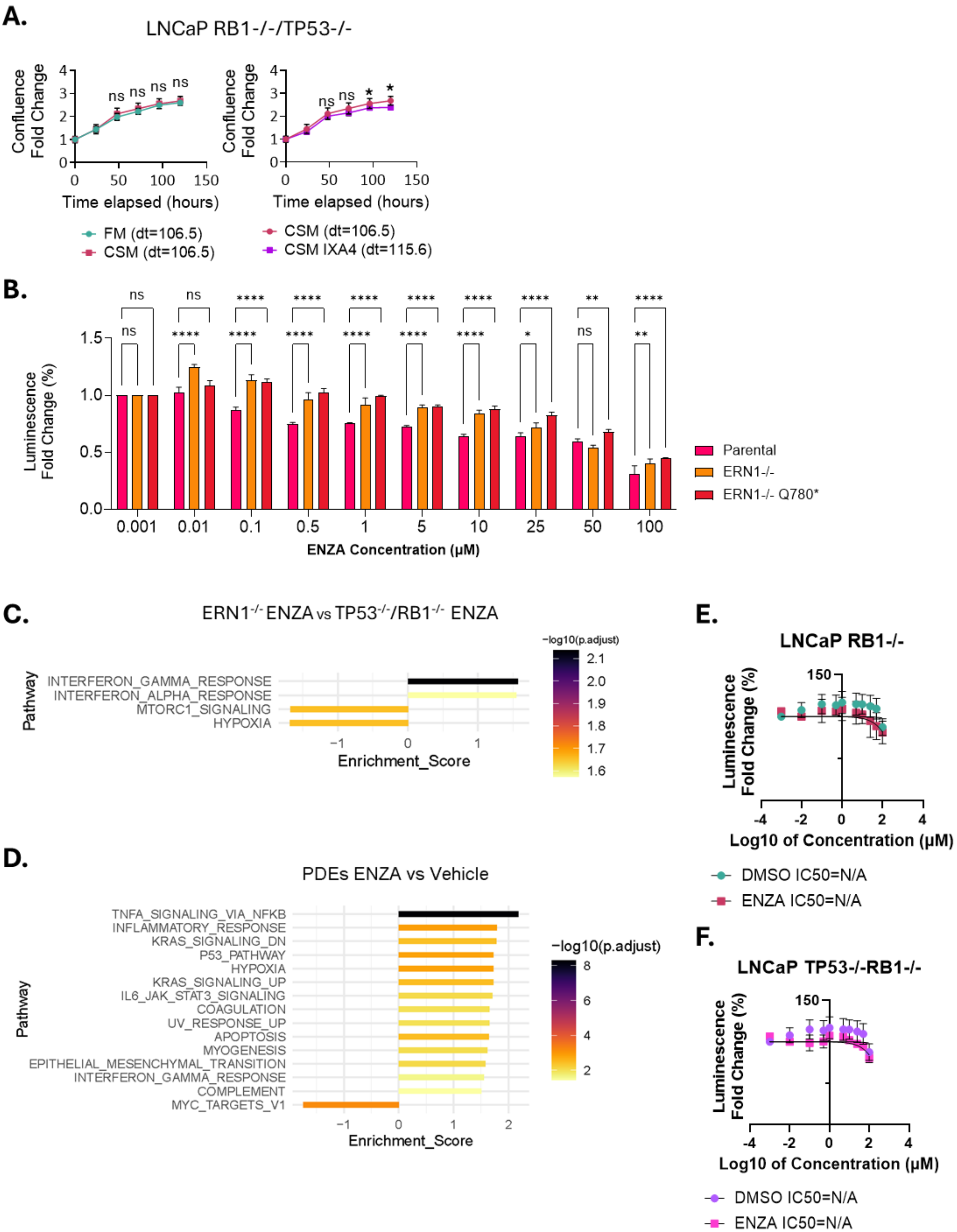

**Supplementary Figure S4**. A) Growth curves of LNCaP RB1$^{-/-}$TP53$^{-/-}$ cells grown over 120 hours in FM (green), CSM (red) or in CSM in the presence of subtoxic (25µM) doses of IXA4 (purple). Confluence fold change was normalised to Incucyte readings taken at the 0-hour time point of each condition. T-tests were used for individual time point comparisons between FM and CSM; CSM plus DMSO and CSM plus IXA4. Exponential (Malthusian) growth curve equations were used to estimate the doubling time of each condition. B) Cell Titer Glo signal fold change in LNCaP, LNCaP ERN1$^{-/-}$ and LNCaP ERN1$^{-/-}$ Q780* cells treated with increasing doses of ENZA over 12 hours. All values are normalised to equivalent DMSO controls for each condition. Statistical analysis was performed using an ordinary one-way ANOVA with Dunnett's multiple comparisons test and single pooled variance. C) Significant Hallmarks upregulated in the RB1$^{-/-}$TP53$^{-/-}$ cells treated with ENZA compared to ERN1$^{-/-}$ cells treated with ENZA. Whole transcriptome RNAseq GSEA Hallmark analysis carried out with bar length signifying Hallmark enrichment score. Bar colour signifies log fold change of Hallmark enrichment compared to control. D) Significant Hallmarks upregulated in PDEs treated with ENZA compared to PDEs treated with vehicle control. Whole transcriptome RNAseq GSEA Hallmark analysis carried out with bar length signifying Hallmark enrichment score. Bar colour signifies log fold change of Hallmark enrichment compared to control. E, F) ENZA concentration gradient measuring cytotoxicity through Cell Titer Glo assays in LNCaP RB1$^{-/-}$ and LNCAP LNCaP RB1$^{-/-}$TP53$^{-/-}$ cells treated for 12 hours with increasing concentrations of ENZA. Logarithmic concentration vs normalised response curves were fit to calculate the IC50 of ENZA for each cell line. Each experiment was performed at least in duplicate. Four stars signify adjusted p value of <0.0001.

Supplementary figure S5

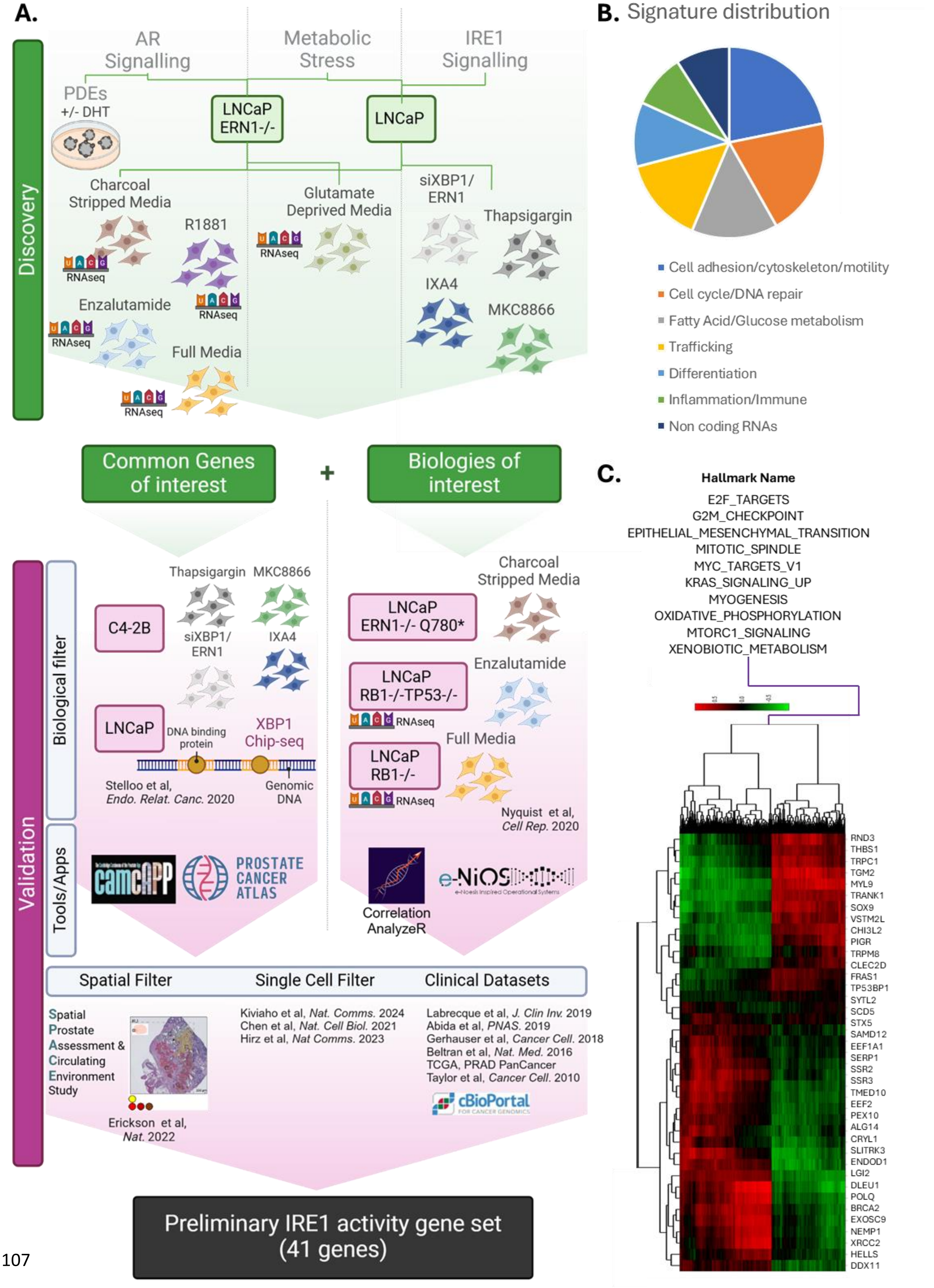

**Supplementary Figure S5**. A) Schematic showing the –omics datasets involved in generating the IRE1 activity gene set. Green lines signify internally generated datasets forming the Discovery component. External and internal datasets used to filter the signature forming the Validation component are highlighted in pink. B) Pie chart indicating the biologies represented within the new IRE1 gene set. These biologies are the result of a GSEA analysis of the final gene list. Created using Biorender.com. C) Heatmap correlating expression of the 41 genes in the IRE1 activity gene set with 500 genes significantly differentially expressed in prostate cancer. The hallmarks these 500 genes represent are displayed above the heatmap. This heatmap was generated using the Correlation AnalyzeR shiny app with the input being the list of 41 IRE1 activity genes, in the topology analysis suite and selecting the tissue type as prostate and sample type as cancer.

#### Supplementary Figure S6

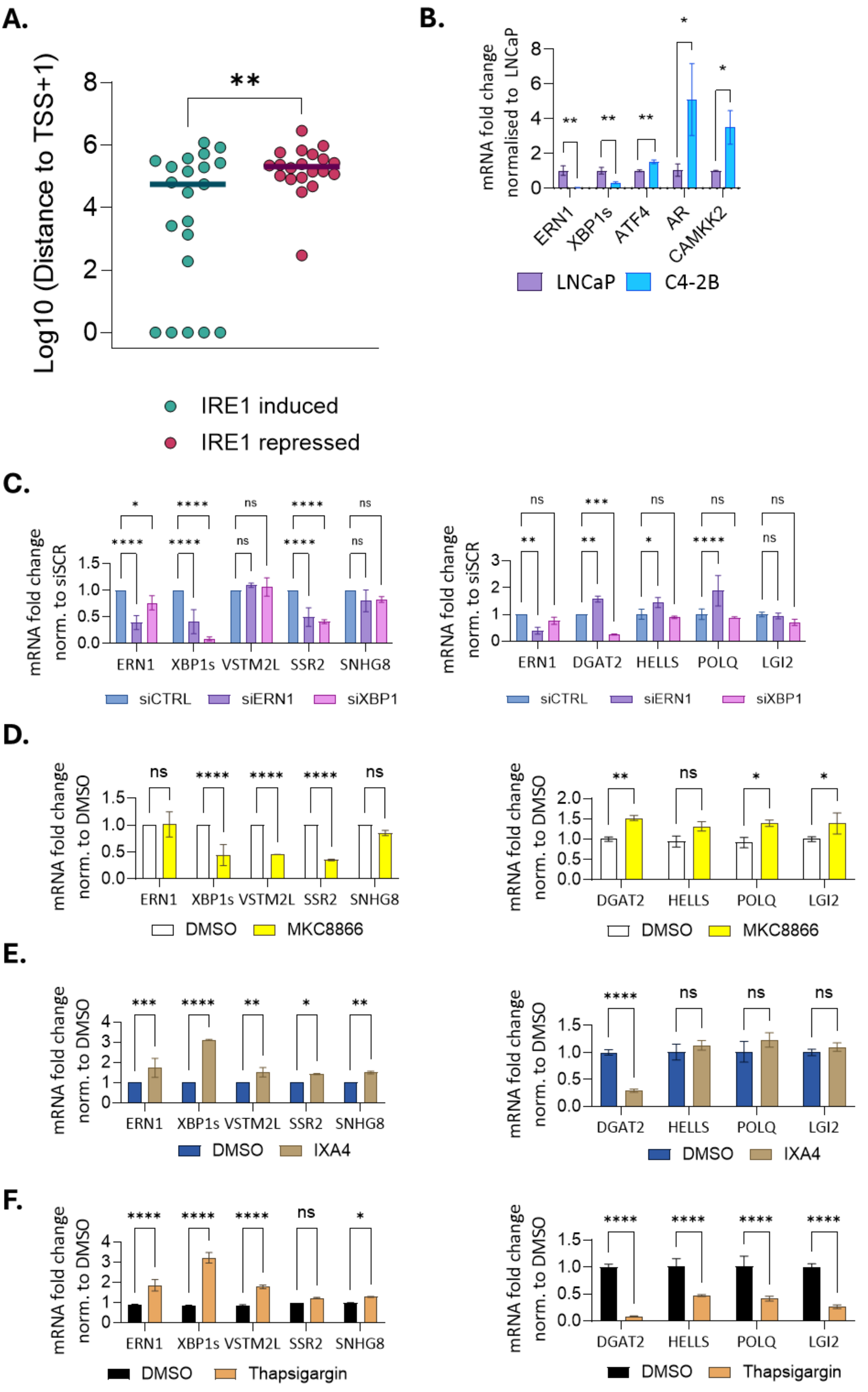

**Supplementary Figure S6.** A) Log10 of the distance between the transcription start site (TSS) of each of the IRE1_41 genes and the nearest significant XBP1 peak. 43% of “IRE1 induced” genes have a an XBP1 peak in their promoter. B) mRNA fold change of *ERN1, XBP1s, ATF4*, full length *AR* and *CAMKK2* in C4-2B cells grown in FM normalised to LNCaP grown in the same FM conditions. Statistical analysis was performed using a T-test per condition. mRNA fold change of *ERN1, XBP1s, DGAT2, VSTM2L, SSR2, SNHG8, HELLS, LGI2* in C42B parental cells treated with C) siRNA against *ERN1* or *XBP1* for 48 hours normalised to scramble control, D) 5µM MKC8866 for 5 hours normalised to DMSO, E) 25µM IXA4 for 5 hours normalised to DMSO and F) 100nM Thapsigargin for 5 hours normalised to DMSO. Statistical analysis was performed using an ordinary two-way ANOVA and Sidak’s multiple comparisons test with single pooled variance. Each experiment was performed at least in duplicate. Four stars signify adjusted p value of <0.0001.

#### Supplementary figure S7

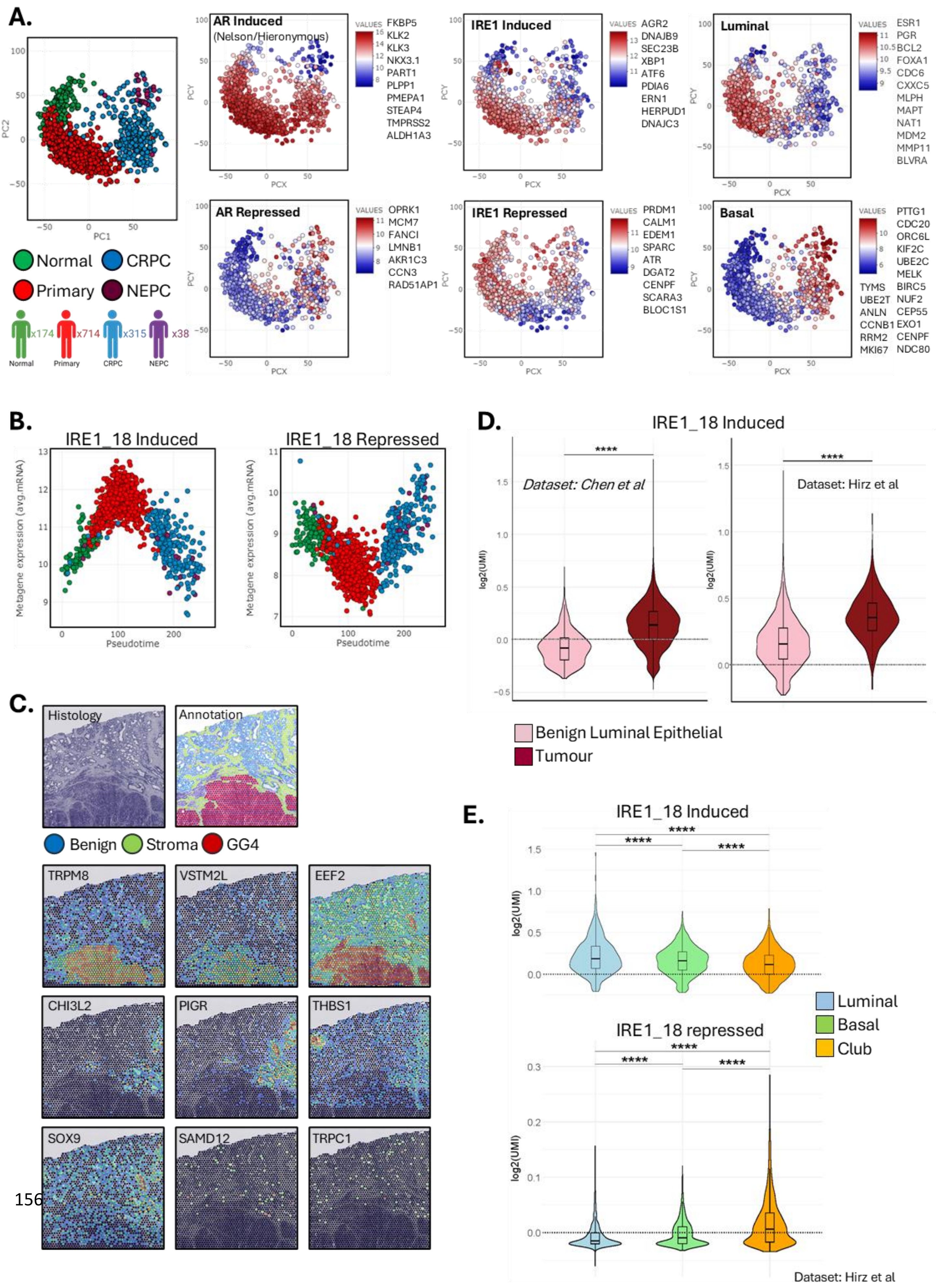

**Supplementary Figure S7**. A) AR, IRE1 and basal/epithelial transcriptomic signatures in a comprehensive prostate cancer transcriptome atlas of integrated RNA-seq datasets from over 1000 clinical tissue samples ranging from benign (green) to NEPC (purple). Heatmaps show distribution of signal (low/blue; high/red) of these signatures across the dataset. B) Unbiased trajectory analysis by inferred pseudotime progression. Plot representing the correlation between mRNAs of IRE1_18 induced and repressed gene set and pseudotime inferred along the trajectory showing the change of these mRNAs across different clinical designations including normal tissue (green), primary tumour (red), AR positive CRPC (blue) and NEPC (purple). C) Example distribution maps of genes within the IRE1_18 gene set representing GG4 tumour, generalised benign or specific benign histological annotations. D) Violin plots showing the enrichment of the IRE1 induced part of the IRE1_18 signature in tumour cells compared to normal epithelial cells in two separate single cell datasets. E) Violin plots showing the enrichment of IRE1_18 induced or IRE1_18 repressed genes, in basal cells and club cells compared to luminal epithelial cells in a single cell dataset.

### Supplementary figure S8

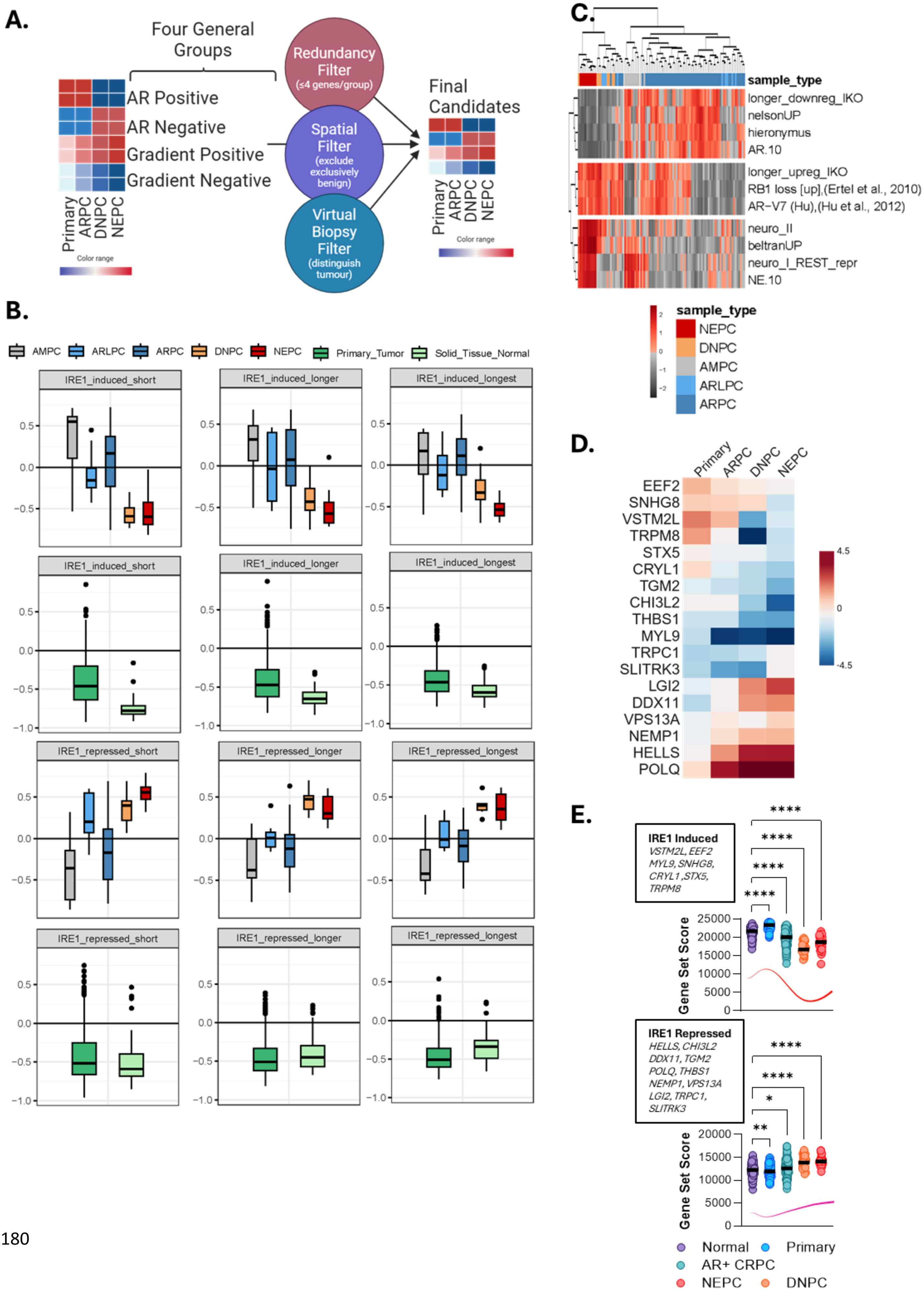

**Supplementary Figure S8**. A) Schematic of filtering and reduction process to produce IRE1_18. Patterns identified in Figure 5F are filtered for redundancy, spatial distribution and bulk prognostication to produce a final list of candidates. Created using Biorender.com B) Box plots of IRE1 activity induced and repressed gene sets, benchmarked in mCRPC (Labreque) and localised disease (TCGA) datasets. Patient populations represented include AR positive (dark blue), AR low (light blue), amphicrine (grey), NEPC (red), double negative (DNPC, orange), normal tissue (light green) and primary tumour (dark green). Using the filtering process described in figure S8A three different lengths of the IRE1 gene set are tested. C) Heatmap of GSVA (single-sample gene set) enrichment scores for selected gene signatures shown across samples from the Labrecque patient cohort (N=98) together with LNCaP parental and ERN1$^{-/-}$ cell line samples (N=5). Colour scale represents regulon enrichment GSVA scores (from lowest in greyscale to highest in red-scale). D) IRE1_18. 18 genes representing different biologies, cell types and clinical designations. Created using Biorender.com. E) Gene set enrichment scores of the IRE1 induced and IRE1 repressed parts of IRE1_18 in normal, primary, AR+ CRPC, DNPC and NEPC samples.

Supplementary figure S9

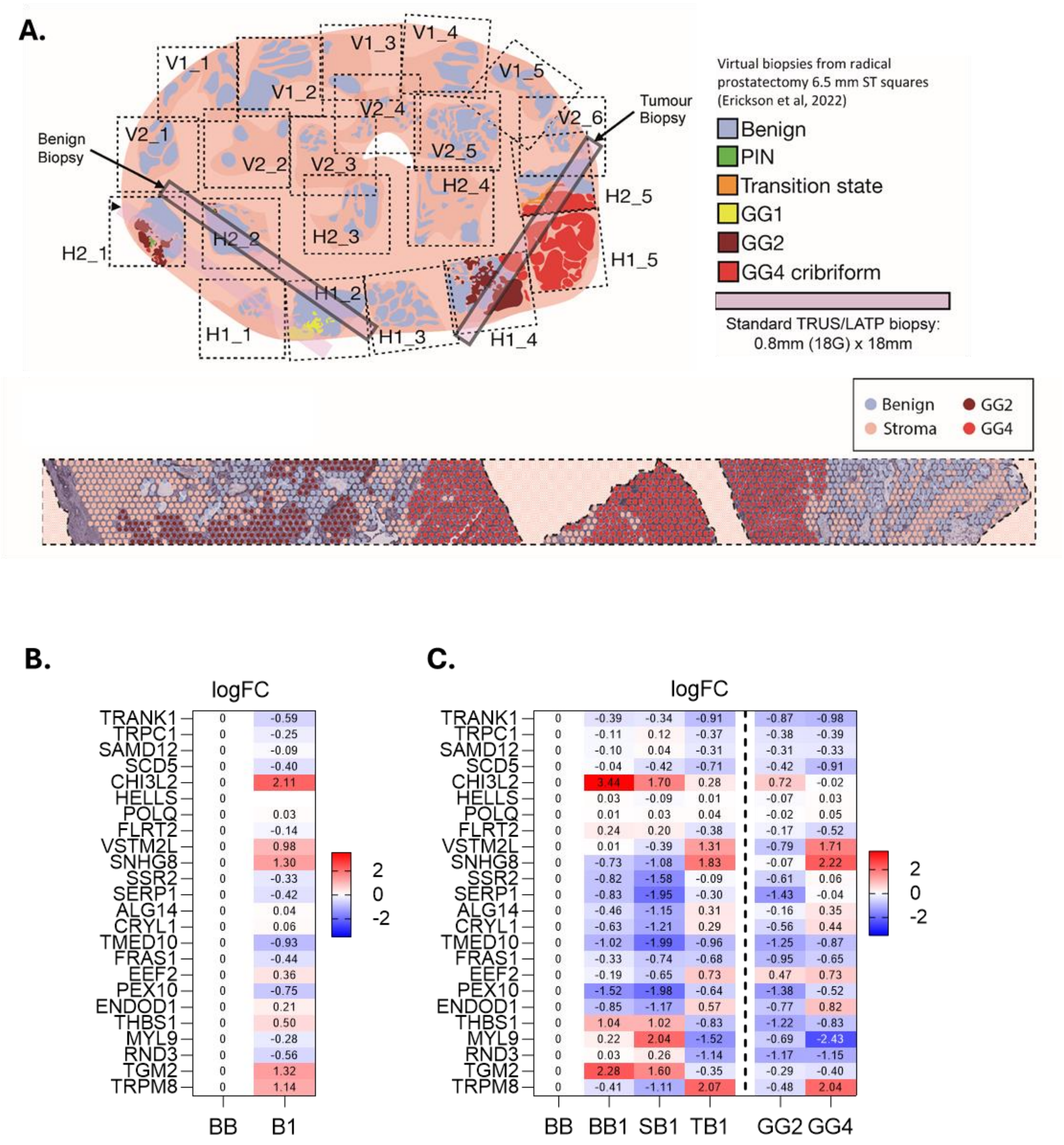

**Supplementary Figure S9**. A) Schematic representing the placement of virtual biopsy containing GG2 and GG4 tumour or benign only tissue (black rectangles). B) Pseudobulk analysis of tumour bearing biopsy (B1) versus benign biopsy (BB). C) Heatmaps showing changes in each sub-compartment within the virtual biopsy compared to the benign biopsy. These comparisons include benign tissue within the biopsy (BB1), stroma (SB1), tumour (TB1) as well as individual grade groups GG2 or GG4). Blue colour signifies downregulation, red colour signifies upregulation. The log2fc values for each gene are represented on the heatmap.
